# Reduced dosage of multiple dynein arm preassembly factors limits cilia regeneration in *Chlamydomonas reinhardtii*

**DOI:** 10.1101/2023.10.31.564885

**Authors:** Gervette M. Penny, Susan K. Dutcher

## Abstract

Motile cilia are complex organelles comprised of >800 structural proteins. Variants in 58 of these genes cause primary ciliary dyskinesia (PCD) in humans. We used a second-site non-complementation approach in diploid *Chlamydomonas reinhardtii* strains to investigate whether reduced dosage of different combinations of motile cilia proteins affects cilia growth / regeneration. Temperature-sensitive mutants in intraflagellar transport genes fail to complement in heterozygous diploids at the restrictive temperature, which may be due to poisonous interactions or a reduction in protein levels. Diploid strains heterozygous for 211 double mutant combinations in 21 genes are phenotypically recessive when unperturbed and do not show SSNC. Consequently, we developed a sensitized screen. When protein synthesis is inhibited, cells utilize the existing cytoplasmic pool of proteins, which is insufficient to build full-length cilia. Six double heterozygous strains in dynein arm preassembly factors (DNAAFs) *pf13, pf23, wdr92,* and *oda8* regrow cilia that are shorter than wild-type diploids, which suggests that double heterozygosity limits the pool of assembled dynein arms further. We isolated a new null allele (*pf23-4)* that shows a more severe loss of ODAs and IDAs in cilia than the previously studied *pf23-1* hypomorphic allele. We also identified a new role for PF23 in cytoplasmic modification of IC138, protein of the I1/*f* inner dynein arm. In our SSNC assay, the *pf23-4* allele also exhibits a more severe phenotype than *pf23-1*. Whole cell extract immunoblots show a reduction of PF23 protein to one-half of wild-type levels in both single and double heterozygous genotypes. The *pf13* mutant shows SSNC with mutants in two outer dynein arm structural proteins, ODA6*, and* ODA9. We suggest that reduction of multiple dynein preassembly factors limits the pool of assembled dynein arms needed for cilia assembly and regeneration. Our data support PF23 as a scaffolding hub protein involved in dynein arm assembly.

**Author Summary:** Motile cilia are essential for movement of cells and fluids. In humans, motile cilia defects cause primary ciliary dyskinesia (PCD), a rare disease characterized by recurring respiratory infections, left-right asymmetry defects, ear infections, and infertility. Many genes are involved in motile cilia function and assembly. We use *Chlamydomonas reinhardtii* to study the effects of changes in protein levels in different combinations of proteins needed for motile cilia. Our results show that the proteins that help fold and assemble the dynein motors in the cytoplasm are sensitive to changes in the dosage of these preassembly factors. Reductions in proteins involved in multiple steps of the pathway prevents efficient dynein preassembly and cilia regeneration under stressful cellular conditions.

## Introduction

Motile cilia are microtubule-based organelles that generate fluid flow or provide cellular motility [1]. In humans, pathogenic variants in more than 58 genes cause a rare disease called primary ciliary dyskinesia (PCD), characterized by recurrent lung infections, neonatal respiratory distress, bronchiectasis, otitis media, situs inversus/ambiguus, and male infertility [2–6]. *Chlamydomonas reinhardtii*, a haploid, single-celled, photosynthetic alga, has been used extensively for the study of motile cilia due to its ease of genetic, biochemical, and microscopic analyses. *C. reinhardtii* has two cilia that are 10-12 µm long that it uses to swim. The cytoskeleton of the cilia (or axoneme) is composed of 9 outer microtubule doublets (MTDs) and two single microtubules known as the central apparatus. A unique feature of *C. reinhardtii* is that when the cilia are removed by pH shock, they will regenerate to full length within 90 min [7]. Rosenbaum and colleagues showed that cilia regeneration requires both the pool of proteins that exist in the cytoplasm after deciliation, and *de novo* protein synthesis [7]. This *de novo* synthesis occurs following a transcriptional upregulation of motile cilia genes begins immediately after deciliation [8,9]. When new protein synthesis is blocked by the addition of cycloheximide, a protein synthesis inhibitor, wild-type haploid cells only regenerate half-length cilia. When this experiment was repeated using *pf16,* a central apparatus mutant, the regenerated cilia were only about 2µm long [7]. In another study, cryo-electron tomography of the *pf16* mutant revealed that in addition to missing PF16, several other central apparatus proteins were missing from the *pf16* mutant cilia [10–12]. These studies highlight two important points. First, the total quantity of proteins available in the cytoplasm is a limiting factor in cilia regeneration. Secondly, the loss of one component necessary for cilia assembly can affect other components in the pool.

The dynein arms are essential for cilia motility. The outer dynein arms (ODAs) are attached to the ciliary axoneme every 24 nm. They generate the force that leads to ciliary bending through sliding and determine the beat frequency of the cilia [13–15]. In *Chlamydomonas,* ODAs are large megadalton complexes comprised of three ATPase heavy chains (α-HC, β-HC, and γ-HC) [16–19], two heterodimer-forming intermediate chains (IC1 and IC2) [20–23], and eleven light chains (LC1-6, LC7a/b, and LC8-10) [24]. The inner dynein arm motors (IDAs) create the asymmetric breaststroke that results in forward swimming [25]. There are 7 IDAs withing a 96 nm repeat. IDA I1/f has two heavy chains, 1α and 1β, intermediate chains IC138, IC140, and IC97, several light chains, and FAP120. The remaining six IDAs (a, c, d, b, e, and g) have one heavy chain and various arrangements of actin, p28, p44, p38, and centrin [26]. ODAs and IDAs are present on all 9 doublets, except MTD1 that lacks ODAs [27,28]. Therefore, to build two full-length functional cilia, approximately 24,000 ODA and 16,000 IDA heavy chains (assuming a cilia length of 12 µm) must be translated, folded and assembled with the other dynein subunits [29]. Furthermore, the ODA heavy chains are over 4000 amino acids long and take approximately 15 min to translate. Since a full-length cilium can be assembled in approximately 90 min, this implies that there is a large biosynthetic demand placed on the cell that needs to be fulfilled in a short period of time [30]. Loss of ODA, IDA and other axonemal components, and the factors needed to assemble or transport them results in the formation of dysfunctional cilia, or the absence of cilia in *Chlamydomonas*. Therefore, we reasoned that it is possible that a change in dosage of multiple ODA or IDA components may hinder the cell’s ability to form sufficient functional dynein motor complexes, and therefore limit cilia regeneration.

Genetic analyses identified proteins required for the proper assembly of ODAs and IDAs known as dynein arm preassembly factors, or DNAAFs [31–33]. DNAAFs localize primarily to the cytoplasm, although there are a few exceptions [34–39]. DNAAFs are a group of structurally diverse proteins that interact with chaperone complexes to guide the folding and stabilization of ODAs and IDAs. They have many different domains that include ones involved in protein-protein interactions and scaffolding functions. For example, PF23 contains several TPR repeats that are well known for their scaffolding functions in other proteins [40,41]. Yeast two-hybrid assays and co-immunoprecipitation of recombinant proteins show that PF23 forms a large scaffolding hub with other preassembly factors that include MOT48, RPAP3, WDR92, FBB18, and ODA7. RUVBL1/2 [42], a member of the R2TP co-chaperone complex [43] also interacts with this complex. Co-immunoprecipitation human cell lysates show that PF13 (Ktu/DNAAF2), a PIH-domain-containing protein also interacts with PF23 (DYX1C1/DNAAF4) [44]. In addition, PF13 forms part of a non-canonical R2TP (Rvb1p- Rvb2p AAA-ATPases, Tah1p, and Pih1p) complex, which is involved in many cellular processes [45,46]. In addition to interacting with several DNAAFs, the WDR (tryptophan-aspartic-repeat) protein WDR92 (Monad) is a part of multiple configurations of co-chaperone complexes including R2TP and prefoldins [46,47]. A different group of proteins involved in dynein assembly are called ‘maturation’ factors. One of the maturation factors, ODA8, appears to be important for efficient preassembly of ODA heavy chains and preparing dyneins to be axoneme-binding competent [48,49]. The molecular events involved in dynein maturation are unclear. Other groups of proteins important for ODA and IDA transport and attachment to the ciliary axoneme include the intraflagellar transport (IFT) trains, the ODA and IDA adapters, and the ODA docking complex [50–54].

The number and diversity of DNAAFs suggests a complex, non-linear role in dynein preassembly. The order of ODA and IDA preassembly and the role that each DNAAF plays in dynein preassembly remains largely unknown. Importantly, existing data outlines the co-operativity of DNAAFs and chaperone protein complexes throughout the dynein preassembly process. In *Chlamydomonas*, loss of DNAAFs results in short and/or paralyzed cilia that lack ODAs and IDAs. In the cytoplasm of some DNAAF mutants, the stability and abundance [33,55–57] of some dynein heavy chains is also affected. These observations suggest that proper folding and stabilization of ODAs and IDAs is necessary to supply the cytoplasmic pool with sufficient dynein components for cilia assembly.

One method to study gene dosage and identify protein products that interact or belong to the same pathway is second-site non-complementation (SSNC). The complementation test is based on the premise that two recessive mutant alleles belong to different genes if they produce a wild-type phenotype when each allele is in *trans* (on different parental chromosomes) [58].If the two heterozygous mutant alleles produce a mutant phenotype when in *trans*, they exhibit a failure to complement and are presumed to belong to the same gene. SSNC occurs when there is failure to complement when the alleles are in different genes [59]. This can occur when both genes involved produce a poisonous protein product. A well-known example involves a cold-sensitive α-tubulin (*tub1-1*) mutation that fails to complement a mutation in β-tubulin (*tub2*) in *Saccharomyces cerevisiae* [60]. Another mechanism of SSNC is through gene dosage. In this model, both genes are heterozygous for null alleles. Therefore, the cell only has half the amount of each gene product. In mice, the *hoxb-5* and *hoxb-6* genes are involved in body segmentation. The double heterozygous mice show skeletal abnormality phenotypes not seen in the single heterozygotes [61].

We utilized well-characterized haploid *Chlamydomonas* strains from the *Chlamydomonas* Resource Center [62,63]. These strains originated from multiple large screens for mutants that show slow-swimming or paralysis [14,28,64]. We used these mutants to generate diploid strains. They were deciliated and allowed to regenerate in cycloheximide. Because de novo protein synthesis is inhibited, the cell is forced to utilize the existing protein pool. This sensitizes the cells to further changes in the dosage of proteins cytoplasmic protein pool. We found that structural genes like ODA and IDA heavy chains generally do not respond to changes in dosage and have no effect on cilia regeneration. In contrast, diploids heterozygous for DNAAFs show sensitivity to reduced quantities of these proteins and regenerate cilia that are shorter than wild-type. In addition, multiple SSNC combinations involve the preassembly factor PF23. Our methodology supports the idea that PF23 is a hub protein and acts at multiple steps in the dynein preassembly pathway.

## Results

### Temperature-sensitive IFT *Chlamydomonas* mutants shows SSNC at restrictive temperatures

IFT trains move proteins into the cilia via the IFTB complex with 16 proteins. Retrograde movement out of the cilia requires the IFTA complex with 6 proteins [65]. Temperature-sensitive mutants of IFT assemble cilia at the permissive temperature (21°C). However, ciliary assembly fails at the restrictive temperature (32°C). Previous work by our group identified mutants in IFT139 and IFT144, (*fla15* and *fla17* respectively) fail to complement in heterozygous diploids at the permissive temperature [66].

We tested additional mutants (see Table 1) to ask whether SSNC occurs between other IFT mutants in IFTA, IFTB, or between IFTA and IFTB complex proteins. We generated double heterozygous diploids of IFTA and IFTB mutants using the methods described in Iomini *et. al* [66] and looked for combinations that fail to assemble cilia. Double heterozygous diploids in four IFTB mutants show SSNC. Two of these are temperature-sensitive (IFT172 and IFT81) and two show an aciliate phenotype at all temperatures (IFT80 and IFT52). Interestingly, *ift72* is the only IFTB mutant that fails to complement IFTA mutants (Fig 1). Among the mutants tested, 5 IFTB mutants fail to complement each other, and 3 IFTA mutants fail to complement in heterozygous diploids. The *ift72* mutant, which belong to IFTB, failed to complement 3 IFTA proteins. (Fig 1).

**Fig 1:**
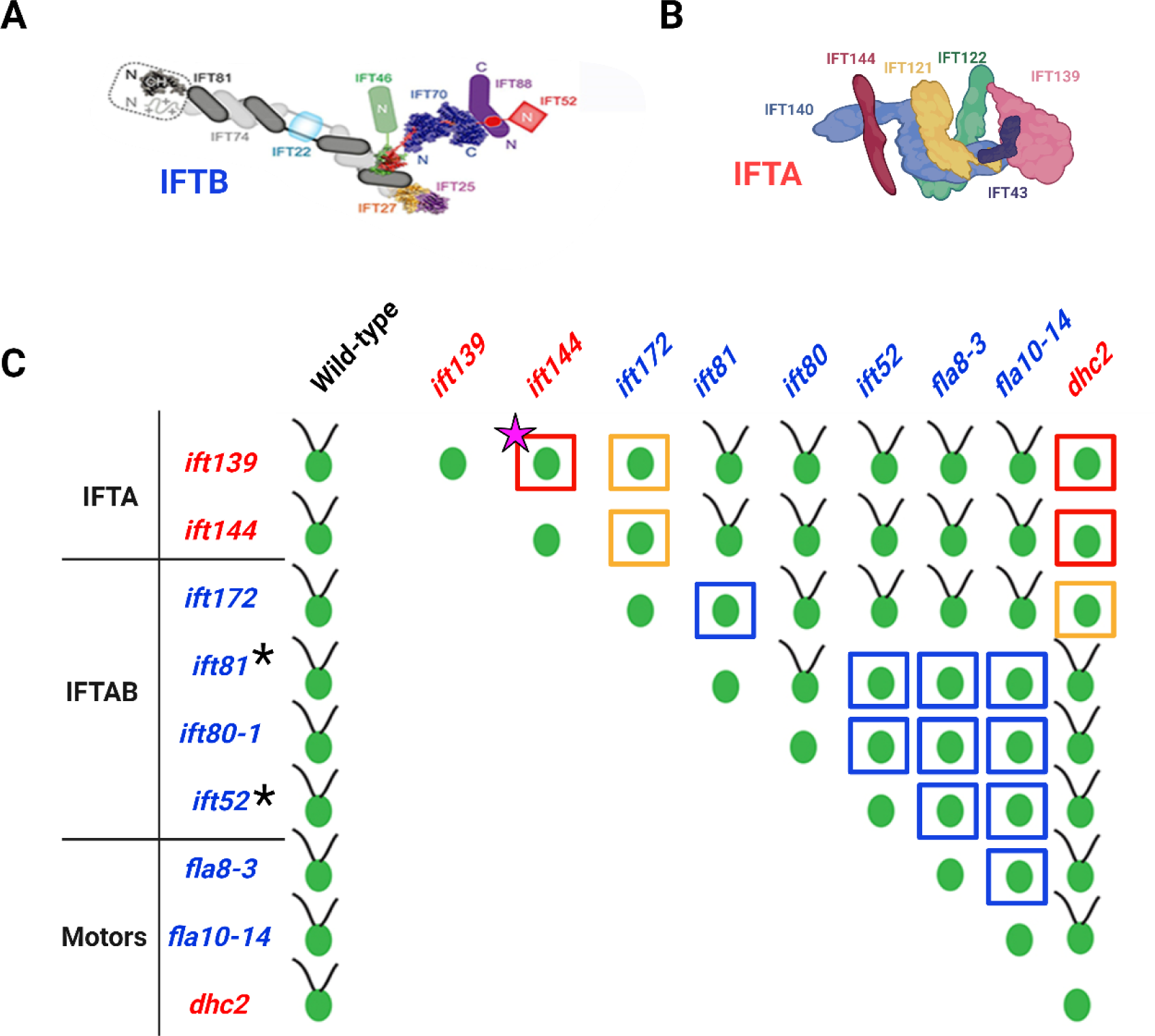
SSNC between double heterozygous mutants in the IFT n *Chlamydomonas* diploid strains. (A) A schematic of the IFTB complex [65]. (B) A schematic of the IFTA complex [65]. (C) The *ift* mutants shown are temperature-sensitive or nonconditional (indicated by *). At the permissive temperature (21^°^C), these mutant strains assemble cilia and swim. After 12 hrs at the restrictive temperature (32^°^C), the strains are aciliate. The mutations are recessive; single heterozygotes assemble cilia at the restrictive temperature (left column). Several members of the same complex fail to complement as double heterozygotes (red squares represent IFTA, blue squares are IFTB). Interestingly, *ift172* complements other components of the IFTB complex but fails to complement mutants of IFTA (yellow squares). The data were generated using methods described in Iomini *et. al.* [66]. The original published SSNC combination of mutants *ift144* and *ift139* is highlighted by a pink star [85].

**Table 1:** Chlamydomonas mutant strains used to generate diploids for SSNC screen. Each mutation is classified by the structural or functional group according to the literature. The human orthologs of each *Chlamydomonas* mutant gene/strain is listed. Dashes indicate no ortholog is found in humans. Mutations in each gene are listed in column along with the known or predicted consequences of the change. Mutations are classed as null or predicted null (PN) or Altered. ^#^Null at the restrictive temperature *\**Sequenced in this work. **Partial function. ^X^CLiP strain [63].

| Category | Mutant | Human Ortholog | PCD Gene | Variant | Type of allele | Ref. |
| --- | --- | --- | --- | --- | --- | --- |
| **Dynein arm assembly factor** | *pf22-1* | *DNAAF3* | Yes | W79X | Null | [69] |
|  | *oda7-1* | *DNAAF1/LRRC50* | Yes | Exon 1 insertion | Null | [70] |
|  | *pf13-1\** | *DNAAF2/KTU* | Yes | 49 kb deletion | Null | [71] |
|  | *pf23-1* | *DNAAF4/DYX1C1* | Yes | In-frame exon 5 deletion | Altered** | [72,73] |
|  | *pf23-4* | *DNAAF4/DYX1C1* | Yes | Insertion exon 1 | Null | This work |
|  | *wdr92-2-295<sup>x</sup>* | *WDR92* | No | Exon 5 deletion, insertion | Null | [57] |
| **Accessory complex** | *oda8* | *LRRC56/MOT37* | Yes | E36fs | PN | [48] |
|  | *oda5* | *CCDC63* | No | V108X | Null | [74,75] |
|  | *oda10* | *CCDC151* | Yes | Unknown | Unknown | [75,76] |
| **ODA docking complex** | *oda1* | *CCDC114* | Yes | E46X | Null | [54] |
|  | *oda3-1\** | *CCDC151* | Yes | M279fs; 1bp deletion | Null | [53] |
|  | *oda14-F28* | None | No | Exon 1 insertion | Null | [77] |
| **ODA intermediate & light chain** | *oda6-95* | *DNAI2* | Yes | V54Gfs | Null | [20,78] |
|  | *oda9\** | *DNAI1* | Yes | K225fs | Null | [79,80] |
|  | *oda12-2* | *DYNLT2* | No | 3'UTR insertion | Null | [81] |
| **ODA heavy chain** | *oda4-1\** | *DNAH9/11/17* | Yes | R2872fs; 5 bp deletion in exon 22 | Null | [80] |
|  | *sup-pf1-1* | *DNAH9/11/17* | Yes | 21bp in-frame deletion in exon 25 | Altered** | [82] |
|  | *oda11-1\** | - | Yes | W778X | Null | [83] |
|  | *oda2-921<sup>x</sup>* | *DNAH5/8* | Yes | Exon 9 insertion | Unknown | [63] |
| **ODA adapter IDA adapter** | *oda16-1* | *WDR69* | No | Insertion unknown | Null | [84] |
|  | *ida3* | None | No | W22X | Null | [85] |
| **Radial spoke** | *pf14* | *RSPH3* | Yes | Q21X | Altered** | [86] |
| **IFTA** | *fla15-1* | *WDR19* | No | C1283R | Altered**/<br>Null# | [66] |
|  | *fla17-1* | *TTC21A* | No | In-frame deletion of 3 exons | Altered**/<br>Null# | [66] |
| **IFTB** | *fla11-1* | *IFT172* | No | L1615P | Altered**/<br>Null# | [87] |
|  | *ift81* | *IFT81* | No | 3' splice site of exon 7 | Altered**/<br>Null# | [88] |
|  | *ift80* | *IFT80* | No | W→X | Null | [89] |
|  | *bld1-1* | *IFT52* | No | Insertion | Null | [90] |
| **IFT Motors** | *fla10-14* | *KIF3A* | No | E24K | Altered**/<br>Null# | [91] |
|  | *fla8-3* | *KIF3B* | No | E24G | Altered**/<br>Null# | [92] |
|  | *fla24-1* | *DYHC2* | No | L3243P | Altered**/<br>Null# | [92] |

### Resequencing of five *Chlamydomonas* mutant strains identifies the causative mutation

A subset of ciliary mutant strains generated by chemical or insertional mutagenesis by the *Chlamydomonas* community was used in our SSNC screen [62,67]. Previously, the causative gene in each mutant strain was identified by rescue with a transgene or by whole genome resequencing. The causative mutations were unknown in five of the strains (*pf13-1, oda3-1*, *oda4-1, oda9*, and *oda11-1*). Therefore, we used whole genome short-read re-sequencing [68]. The mutations in strains *oda3-1, oda9,* and *oda11-1* were identified (Table 1, Supplemental Table 1). However, the SnpEff analysis pipeline did not reveal any candidate mutations in *pf13-1* and *oda4-1*. After manual curation of sequence from the *pf13-1* strain, we identified a 44.7 kb deletion that removes the *PF13* gene and 6 other genes (Table 1, Supplemental Table 1, Supplemental Fig S1A, S1B). PCR shows that the centromere-adjacent breakpoint removes all the exons of *PF13* (exons 3, 7 and 10 were tested), but the 3’UTR is retained. The telomere-adjacent breakpoint involves Cre09.g411700, which contains 3 exons and is the last gene disrupted by the deletion.

Only a small part of the 3’ end of exon 3 is deleted; exons 1, 2 are retained (Supplemental Fig S1). The *pf13-1* mutation was rescued by a transgene generated by recombineering [93,94] containing the wild-type *PF13* gene. This indicates that deletion of *PF13*, but not the other 5 genes, causes the phenotype observed in *pf13-1* (Supplemental Fig S2). The reads at the *ODA4* (*DHC14*) locus were manually examined. We found a 5 base-pair deletion in the *oda4-1* mutant that changes the reading frame in one of the AAA ATPase domain (Supplemental Table 1, Supplemental Fig S1C). PCR analysis confirmed that the sequence changes identified by re-sequencing of *oda3-1, oda4-1, oda9,* and *oda11-1* are present in the mutant strains (Supplemental Fig S1, Supplemental Table 2).

The 22 strains we selected fall into five classes that alter the assembly of the outer and/or inner dynein arms or their transport into cilia (Fig 2A-2E). To verify the phenotypes of the strains, we examined cilia length using immunofluorescence with an antibody to acetylated α-tubulin that stains cilia and rootlet microtubules (Fig 2F). As expected, the *oda* strains assemble full-length cilia. Strains with preassembly factor mutations (*pf22-1*, *pf13-1*, *pf23-1,* and *wdr92-2*) have short, immotile cilia. One known exception is the preassembly factor mutant *oda7* that has full-length cilia and is motile (Fig 2F, 2G, 2H). The swimming phenotype of *oda7* resembles the slow, jerky movements characteristic of *oda* mutants [95–98]. Swimming velocities vary among strains. In general, *oda* strains swim at about one-half the wild-type speed and have a reduced beat frequency [15,99]. Most of the *oda16-1* cells were immotile, but approximately 10% of cells swam slowly. The *ida3* strain swims more slowly than wild-type but has a wild-type beat frequency and wild-type cilia length (Fig 2G, 2H) [52,100].

**Fig 2.**
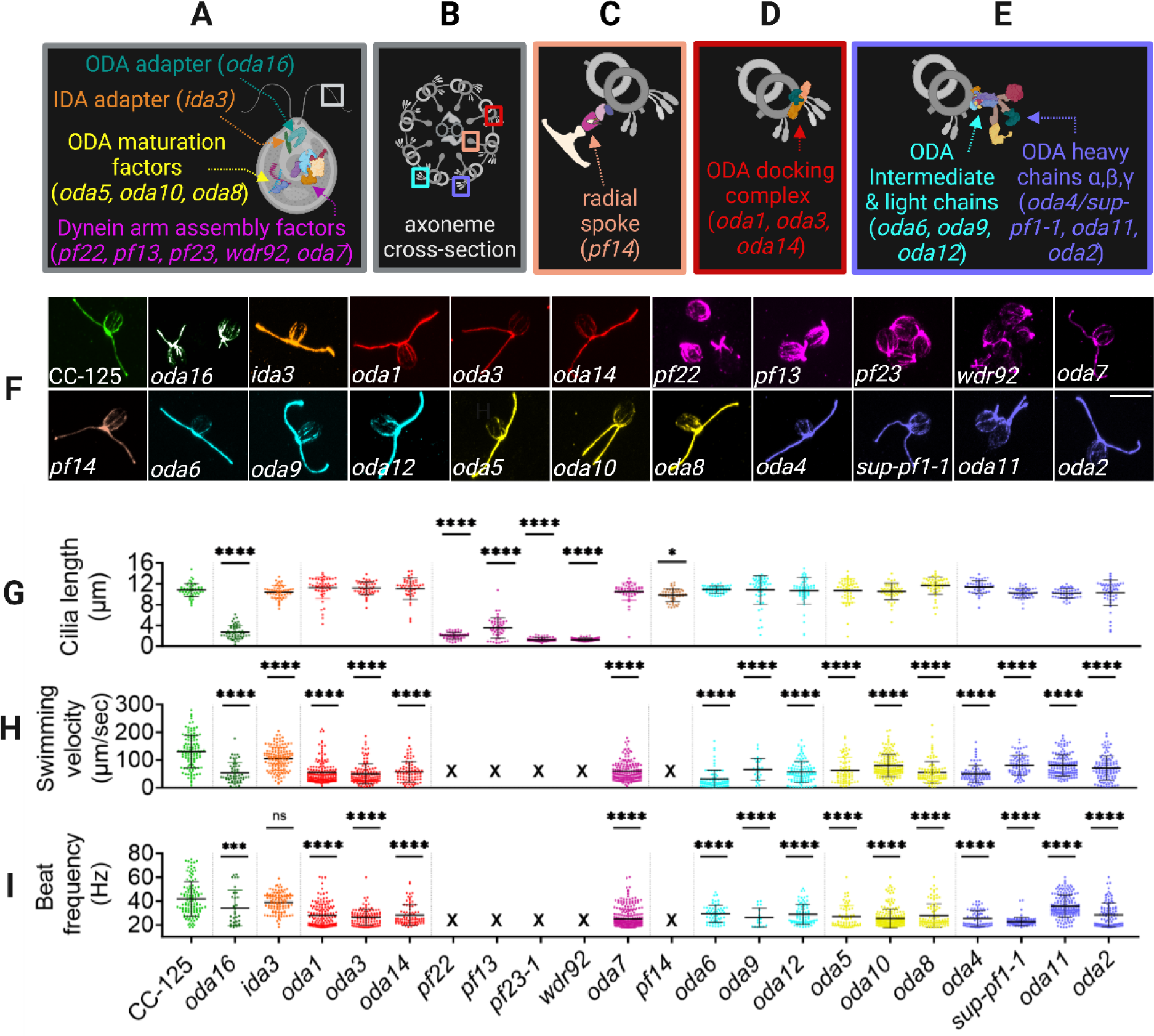
Different classes of motile cilia genes and dynein preassembly factor mutants have diverse phenotypes. (A-E) Motile cilia genes in this study are grouped according to their function or the structure affected. The mutant strains in each group is listed in parentheses. (A) Dynein arm preassembly factors (pink), ODA maturation factors (yellow), ODA adapter (dark green) and IDA adapter (orange). (B) A cross-section of the *Chlamydomonas* cilium outlined by the white box in (A).The boxes highlight the locations of four structural groups; they are the radial spoke (tan, C) the ODA docking complex (red, D), and the ODA intermediate and light chains (cyan, E) and ODA heavy chains (purple, E). (F) Wild-type *Chlamydomonas* and mutants from each functional group in A-E are stained with antibody against acetylated α-tubulin to show the cilia and rootlet microtubules. (G) Quantification of cilia lengths. One cilium from 50 cells were measured for each strain. Statistical analysis was performed using a One-Way ANOVA with adjustments for multiple comparisons. Each strain is compared to wild-type strain CC-125. (H) Swimming velocity measurements for each strain in F. Strains *pf22*, *pf13*, *pf23-1*, *wdr92* and *pf14* are non-motile and are indicated by an ‘X’. (I) Beat frequency measurements for each strain in F. ‘*’ p < 0.05; ‘**’ p < 0.001; ‘*** p < 0.0001; ‘****’ p < 0.00001; ‘ns’ not significant.

### Cilia length is regulated independently of cell size in *Chlamydomonas*

#### To identify candidate gene pairs that show SSNC between motile cilia mutants in *Chlamydomonas,* we generated heterozygous diploid strains. One haploid strain with a mutation in *gene 1* and wild-type for *GENE 2* was mated to a second haploid strain with a mutation in *gene 2* and wild-type for *GENE 1* (Fig 3). Each strain contains a selectable genotype for acetate auxotrophy (*ac17*) or paromomycin resistance (*aphviii*). The *ac17* mutation and *aphviii* selectable marker was obtained by crosses to strains CC-5908 and CC-5909 (see Materials and Methods). The *aphviii* marker was selected in place of *nit2*

1. [66] because of stronger selection. The selectable genotypes are reciprocal in each haploid strain. The outcome of this cross is a diploid that is heterozygous for two ciliary mutants (Fig 3A). Diploids appear as bright green colonies on medium lacking acetate and containing paromomycin (Fig 3B). In addition, to verify the recessive nature of each allele, single heterozygous diploids were generated by a cross between a wild-type haploid strain and each mutant strain.

**Fig 3:**
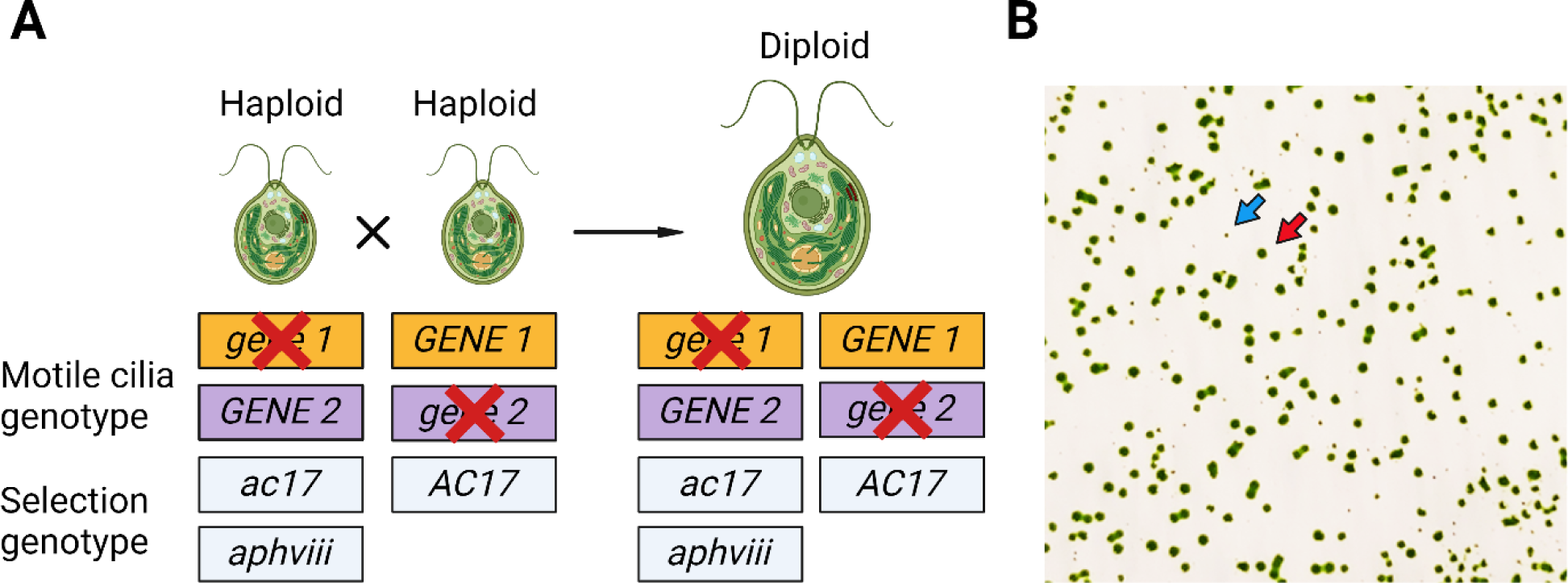
Generation of *Chlamydomonas* diploid strains. (A) Diploids were selected after 3-5 days on medium lacking acetate and containing paromomycin. Haploid parental strains with the genotype *ac17; aphviii* (CC-5908 and CC-5909) die on plates that lack acetate although they are resistant to paromomycin. Haploid parental strains with the genotype *AC17* grow in the absence of acetate and die on medium with paromomycin. Only diploid cells survive on plates lacking acetate and containing paromomycin. The two sets of haploid cells each carry mutations in two different cilia genes (*gene 1* and *gene 2*). Mating these strains produces diploid cells that are heterozygous for both mutations. (B) Colonies from diploid cells are bright green (red arrow) on a pale green background of haploid cells (blue arrow).

We compared haploid and diploid wild-type *Chlamydomonas* vegetative cells. Previous analyses of diploid cells reported differences in cell size, mating-type, and DNA content compared to haploid cells [101]. Mating-type PCR analysis shows that diploid cells contain both mating-type plus and mating-type minus loci [102,103]. Diploids generated in this work were also genotyped using PCR primers for the heterozygous selectable marker genes used to select them, they are *ac17/AC17*; *aphvii*i (Fig 3A, Supplemental Fig S3). Diploid cells appear larger and swim more slowly than haploid cells by phase microscopy. To quantitate the size of diploid cells, we performed immunofluorescence on haploid and diploid wild-type strains. We found that diploid cells are longer (8.5 microns compared to 6.9 microns for haploids) (Fig 4B, 4F, Supplemental Table 4); these data agree with results from Ebersold [101]. Diploid strains exhibit slower swimming velocities and beat frequencies than haploid strains (Fig 4B, 4C, 4D). The cilia of diploid cells are the same length as haploid cells (Fig 4E).

**Fig 4:**
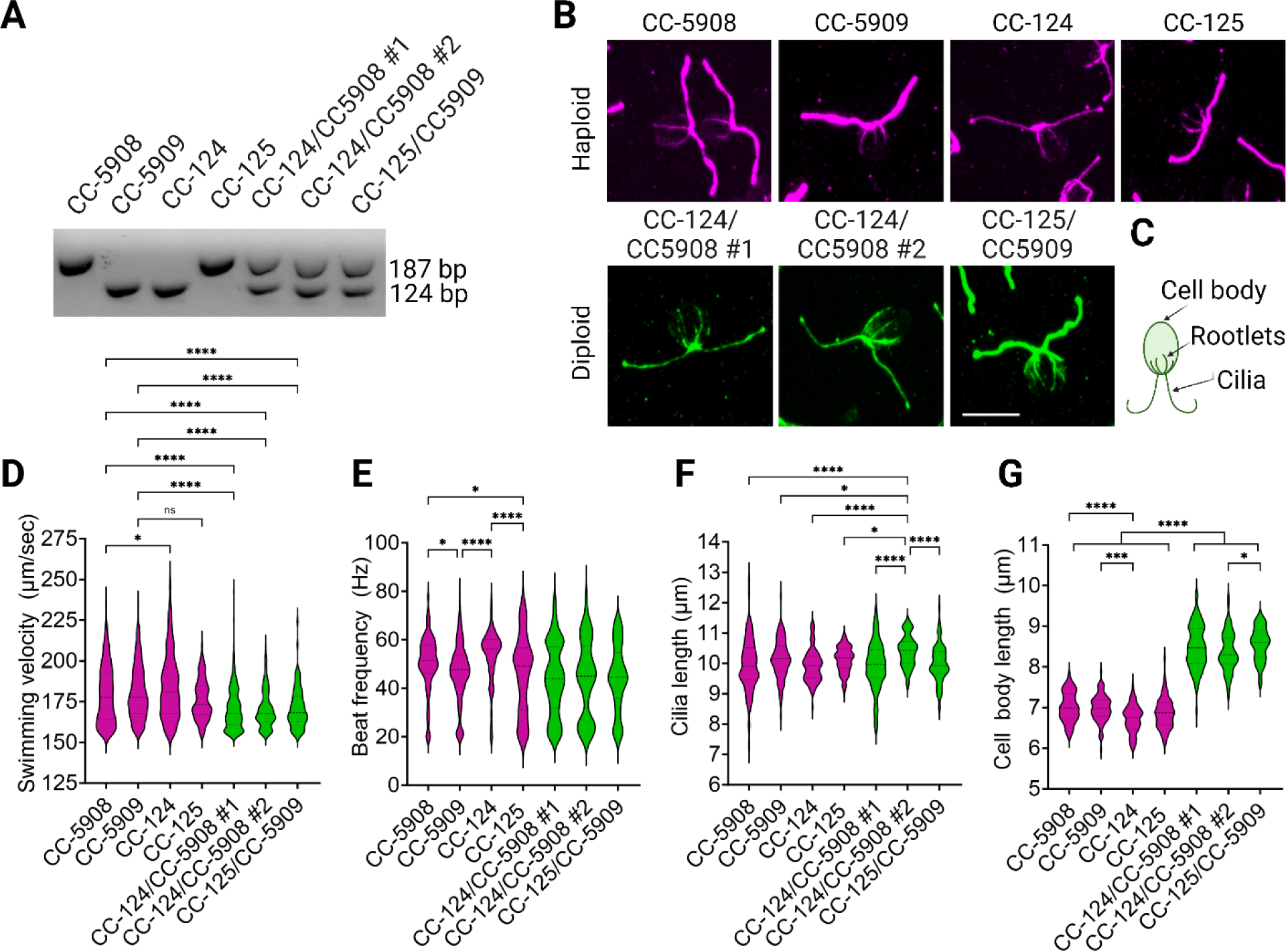
A comparison of haploid and diploid *Chlamydomonas* strains. (A) PCR amplicons for the *MTD* gene (mating-type minus with 124 bp product) and *MTA* gene (mating-type plus with 187 bp product) distinguish haploid and diploid strains. (B) Immunofluorescence of vegetative haploid strains with the genotype *ac17; aphviii (*CC- 5908, CC-5909), CC-124 and CC-125 wild-type strains, and diploid strains generated by crosses between CC-5908 and CC-124 or CC-5909 and CC-125. Cells stained with an antibody against acetylated α-tubulin. (C) An illustration of the microtubules containing acetylated tubulin is provided for reference. (D) Swimming velocity, (E) Beat frequency, (F) Cilia length and (G) Cell body length measured from the apex of the cell to the base of the cilia of strains shown in B. Statistical analysis is a one-way ANOVA using an all-by-all comparison. ‘*’ p < 0.05; ‘**’ p < 0.001; ‘*** p < 0.0001; ‘****’ p < 0.00001; ‘ns’ not significant.

These data suggest that cilia length is regulated independently of cell size and the pool of proteins present in the cell. The strains in Table 1 were used to construct 1 wild-type, 21 single heterozygous, and 211 double heterozygous strains. None of the strains display parental mutant phenotypes. We quantified cilia length, swimming velocity, and cilia beat frequency of a subset of vegetative single and double heterozygous diploids. There were no differences in cilia length, swimming velocity, or beat frequency between wild-type and heterozygous diploids (Fig 5). Additionally, there was no obvious pattern of motility defects that were detected in 211 double heterozygotes.

**Fig 5:**
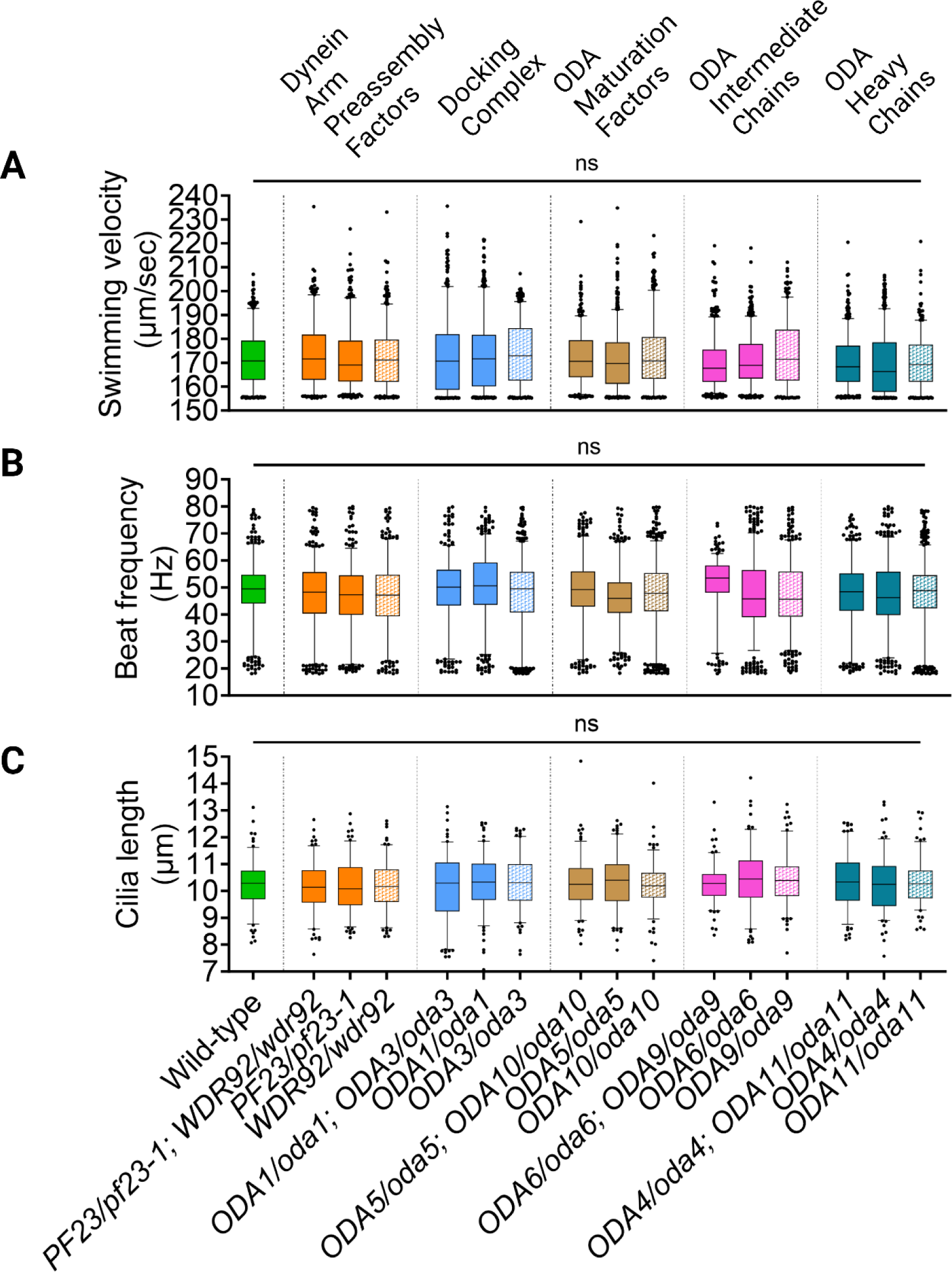
Motile cilia structural and dynein preassembly factor genes are recessive in vegetative diploids. A subset of diploids was examined for the occurrence of SNNC in 5 double heterozygous strains with single heterozygous controls. (A) Swimming velocity, (B) Beat frequency, and (C) Cilia length for wild-type diploids and heterozygous diploids with mutations in different ODA preassembly or structural genes. Pooled data for 3 biological replicates are shown for each strain. N= 50 cilia from 50 cells for each replicate. The individual data points shown are below and above the 10^th^ and 90^th^ percentile. Solid green fill indicates wild-type and single heterozygotes are represented by various other solid fill colors. Striped fill indicates double heterozygotes. Statistical analysis is a one-way ANOVA. ‘*’ p < 0.05; ‘**’ p < 0.001; ‘*** p < 0.0001; ‘****’ p < 0.00001; ‘ns’ not significant.

### Diploid cells have a larger cytoplasmic pool of proteins available to regenerate cilia compared to haploid cells

To develop a sensitized assay to observe changes in cilia length, we relied on results from Rosenbaum, Moulder & Ringo who assessed the effect of cycloheximide on cilia preassembly in haploid cells [7]. They found that wild-type haploid cilia regenerate to approximately 5 to 6 µm in 10 μg/ml cycloheximide following deciliation. The cilia in a *pf16-1*, a central apparatus mutant, regrew cilia that were less than 2 microns. These data showed that the pool of proteins in wild-type cells is only sufficient to build half-length cilia and that the pool is further limited in a mutant *pf16* background. Previous experiments in haploid strains showed that loss of both the inner and outer dynein arms results in short cilia [15,104–106]. SSNC that affects both the ODA and IDAs could show a phenotype. We tested for a defect in cilia length that could be observed in our heterozygous diploids when subjected to this stress condition.

First, we assessed whether wild-type diploid *Chlamydomonas* cells regenerated cilia similarly to haploid strains in the absence of cycloheximide. We included haploid strains *pf16* and *pf16B* as controls. They are renamed *pf16-1* and *pf16-2,* respectively, to agree with the current *Chlamydomonas* nomenclature. The strains were deciliated and allowed to regenerate in the presence or absence of 10 µg/ml cycloheximide (Fig 6A). All wild-type haploid and diploid cells regenerate to their original pre-deciliation length in the absence of cycloheximide. Diploid cells regenerate more quickly than haploid cells under both conditions. At 40 mins post-deciliation, diploid cilia are approximately 8 microns long in the absence (Fig 6A) or 5 microns in the presence (Fig 6B) of cycloheximide while haploid cilia average 6 or 4 microns. The *pf16* strains fail to regenerate to their original length after deciliation; the defect in the *pf16-2* strain is more severe. The *pf16-2/pf16-2* diploids also regenerate longer cilia than the *pf16-2* haploid strains under both conditions (Fig 6B, 6C). Our data recapitulate the observations of Rosenbaum *et. al.* [7]. Cilia length does not change drastically after 180 min of regeneration. Our data are consistent with the hypothesis that the pool of proteins needed for building cilia is larger in diploid cells since they assemble longer cilia and regenerate more quickly.

**Fig 6:**
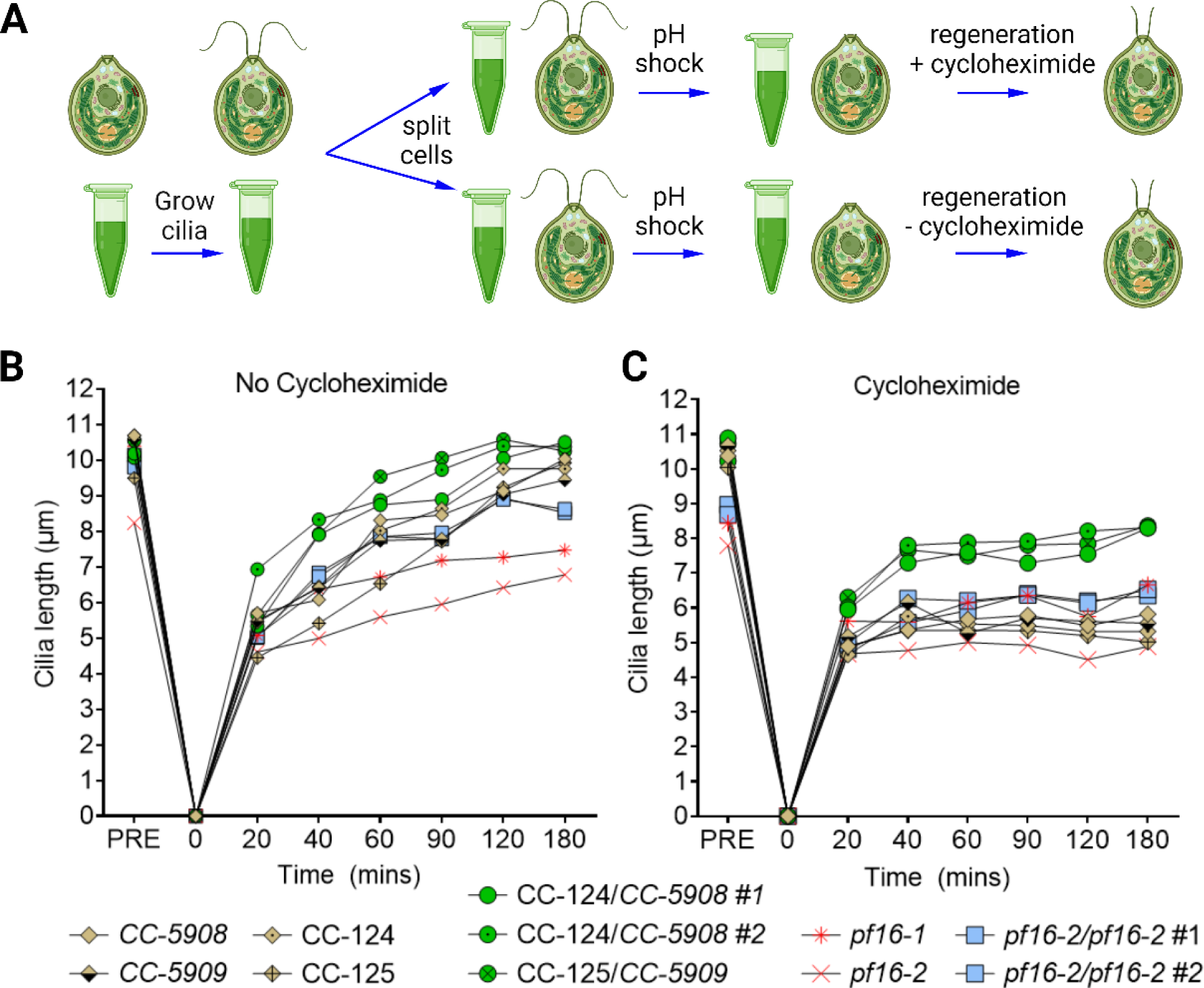
Cycloheximide reduces cilia regeneration after deciliation in haploid and diploid cells to different extents. (A) Vegetative cells with the indicated genotypes were deciliated pH shock and allowed to regenerate for 180 min. Samples were collected for cilia length measurements at the timepoints shown. 100 individual cilia from 100 cells were measured for each strain and timepoint. (B) A parallel experiment was performed with 10 μg/ml cycloheximide added post-deciliation. The behavior of haploid wild-type strains are shown in brown, wild-type diploid cells are green, *pf16-1* and *pf16-2* haploid cells are red, and *pf16-2* homozygous diploid cells are blue.

### Diploids heterozygous for mutations in DNAAFs show SSNC in deciliation screen

Having established that diploid cells follow a similar regeneration pattern to haploid cells in the presence of cycloheximide, we asked if any of the single or double heterozygous strains show a cilia regeneration phenotype. Using the protocol outlined in Fig 6A, we deciliated the single and double heterozygous strains, and wild-type diploids, and allowed them to regenerate in cycloheximide for 40 mins.

As shown in Fig 7 and Supplemental Table 8 (rightmost column), all the mutations in the single heterozygous strains tested in this assay are recessive to the wild-type allele and support previous data from both human and model organism studies (Table 1). Double heterozygous strains with mutations in structural proteins (the radial spoke gene *PF14 (RSP3),* the ODA heavy, intermediate, and light chains (*oda4*, *sup-pf1-1*, *oda11*, *oda2*, *oda6*, *oda9* and *oda12*), and ODA-DC (*oda1*, *oda3*, and *oda14*) all complement. The IFT adapters, *oda16* and *ida3*, as well as the maturation factors *oda5* and *oda10* also complement. These combinations do not show SSNC.

**Fig 7:**
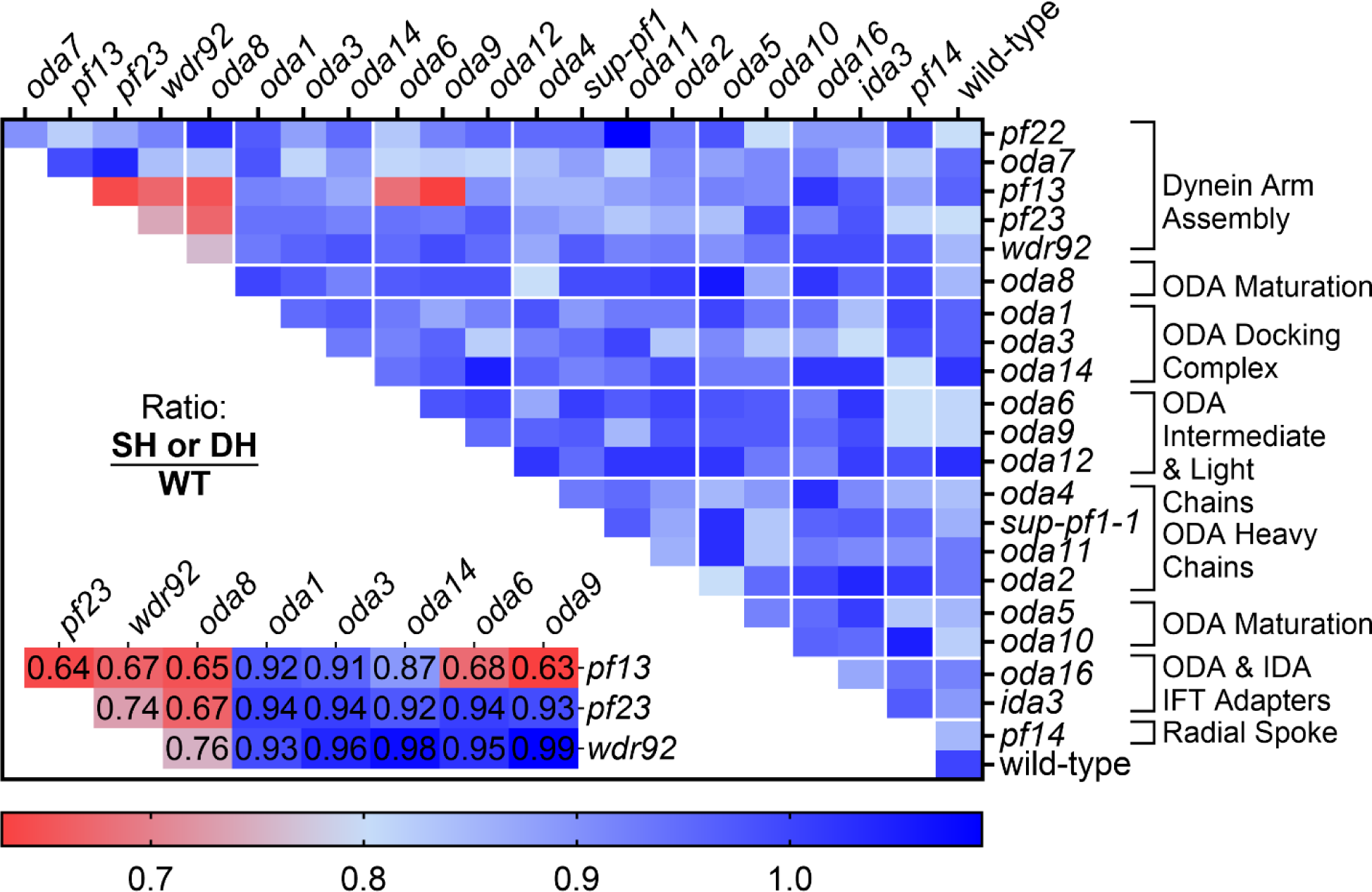
Cilia lengths following regeneration in cycloheximide for heterozygous diploids. Each strain and a wild-type control was deciliated and allowed to regenerate for 40 min in 10 μg/ml cycloheximide. Three biological replicates were measured for each heterozygous diploid. One biological replicate was measured for the wild-type diploid. For each replicate, 100 individual cilia from 100 cells were measured and the average length was calculated. Each box shown in the heat map represents the product of the average length of the pooled biological replicates for each single (SH) (rightmost column) or double heterozygote (DH) divided by the average length of the wild-type diploid (WT). Cooler colors (blue) indicate that the cilia length of the tested heterozygotes is similar to wild-type cilia length. Warmer colors (red) indicate that cilia length is shorter than wild-type cilia length. An inset of the candidate hits in red is shown in the bottom left corner with the cilia length ratio value.

We found a low occurrence of SSNC in our screen, but it is specific. Non-complementation was observed only when at least one of the mutations was in a dynein preassembly factor gene. SSNC was observed between *pf13* and either *oda6* or oda9. A mutation in the dynein maturation factor gene *ODA8* shows SSNC with *wdr92, pf13,* and *pf23-1*. The *pf23* mutation shows SSNC with *pf13-1* and *wdr92* (Fig 7, inset lower left).

#### A null *Chlamydomonas pf23* mutant completely lacks ODAs and IDAs in the cilia

Most of the alleles that show SSNC in our screen are null alleles. The *pf23-1* allele is unusual because it contains an in-frame deletion of exon 5 and surrounding introns that produces a smaller PF23 protein with reduced function [73]. We hypothesized that a null *PF23* allele in *Chlamydomonas* would have greater effects on dynein preassembly and lead to a more severe SSNC phenotype.

We utilized a CRISPR/Cas9 site-directed insertional mutagenesis approach to insert the *aphvii* cassette that confers hygromycin resistance into exon 1 of the *PF23* gene [107] in the wild-type strain CC-5908. We obtained three independent strains from two separate transformations. These three strains were renamed *pf23-2, pf23-3,* and *pf23-4* [107] (Supplemental Fig S5). Immunofluorescence analysis of gametic wild-type and *pf23* mutant strains show that the *pf23-1* strain assembles cilia that are 2.14 +/- 0.86 microns long, while the *pf23* null strains assemble cilia that average 0.87 +/- 0.23 microns in length (Fig 8A). Immunoblot analysis using an anti-PF23 antibody [57] shows that the CRISPR insertional mutants completely lack the PF23 protein, unlike the *pf23-1* strain that produces a smaller PF23 protein (Fig 8B). This indicates that complete loss of PF23 protein results in a more severe phenotype than in *pf23-1* strain.

**Fig 8:**
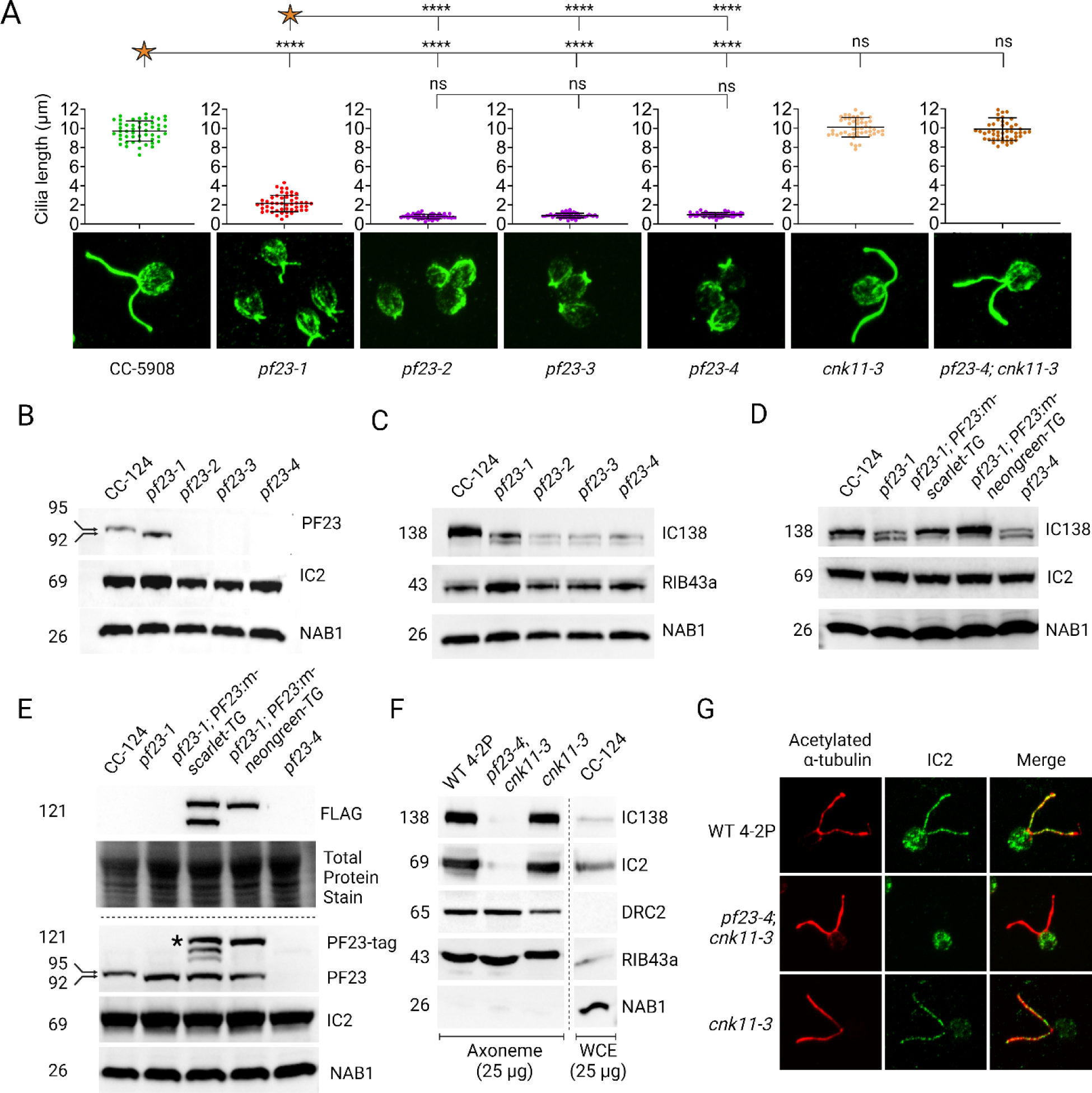
Characterization of null *pf23* alleles generated by CRISPR/Cas9 site-directed insertional mutagenesis. (A) Immunofluorescence staining of gametic cells using antibody to acetylated α-tubulin to visualize cilia length of wild-type, *pf23-1*, and the three CRISPR generated *pf23* strains. The quantification of length for each strain is positioned directly above its corresponding image. (B) Immunoblot probing gametic whole cell extracts for PF23 and IC2 intermediate chain proteins that shows that these alleles do not produce PF23 protein and are null. (C) Immunoblot of *pf23* mutant strains for IC138 I1/f intermediate chain and MIP RIB43a. NAB1 is used as cytoplasmic loading control. (D) Immunoblot of *pf23-1*, *pf23-4,* and two independent transgene strains *pf23- 1; PF23:m-scarlet-TG* and *pf23-1; PF23-neon green-TG* that show rescue of the ciliary preassembly defect. (E) Strains in (D) were probed with the PF23 antibody and anti-FLAG to show rescue of PF23 protein expression. (F) Immunoblot of isolated cilia of wild-type, *pf23-4; cnk11-3*, *cnk11-3,* and CC-124 for various axonemal components. NAB1 is used to assess cytoplasmic contamination. The NAB1 image shown is overexposed to show the relative abundance in CC-124 whole cell extracts compared to its absence in axonemes. (G) Immunofluorescence of strains in (F) using antibodies against IC138 and IC2.

Meiotic analysis of all three strains showed that *pf23-2* and *pf23-3* produced meiotic inviability, while crosses with *pf23-4* produced primarily tetrads with four viable progeny. Long-range sequencing of *pf23-2* and *pf23-3* showed that these strains contain translocations between the *PF23* gene and chromosome 5 and 3, respectively [107]. Because *pf23-4* did not show any meiotic abnormalities, it is unlikely that it possesses a translocation. Therefore, we used strain *pf23-4* for our SSNC studies.

We characterized the effects of the *pf23-4* null allele on dynein arm preassembly using mass spectrometry of isolated axonemes. However, the *pf23-4* cilia are extremely short and difficult to isolate. Mutations in *CNK11,* a NIMA-related kinase, partially suppresses the cilia length defects of other strains without rescue of the motility defects [105,108]. The *pf23-4* short cilia phenotype is also rescued by *cnk11* (Fig 9). The proteomics analysis of *pf23-1; cnk11-3* axonemes showed almost complete loss of the IDA proteins, while ODA species were reduced by approximately 50 to 60 percent [73]. Our analysis reveals that the *pf23-4; cnk11-3* strain results in the complete absence of both ODAs and IDAs from isolated axonemes (Table 2, red shading). Surprisingly, ODA docking complex proteins are moderately reduced by approximately 60% (tan shading). Components of other axonemal proteins including MIA, radial spokes, N-DRC, and the central apparatus remain largely unaffected, or only mildly reduced (green shading). There is a drastic loss of the I1/f tether component FAP44 (tan shading), but the tether-head protein FAP43 is retained. The *cnk11-3* mutation had little effect on the protein composition of the cilia (Supplemental Table 6).

**Fig 9:**
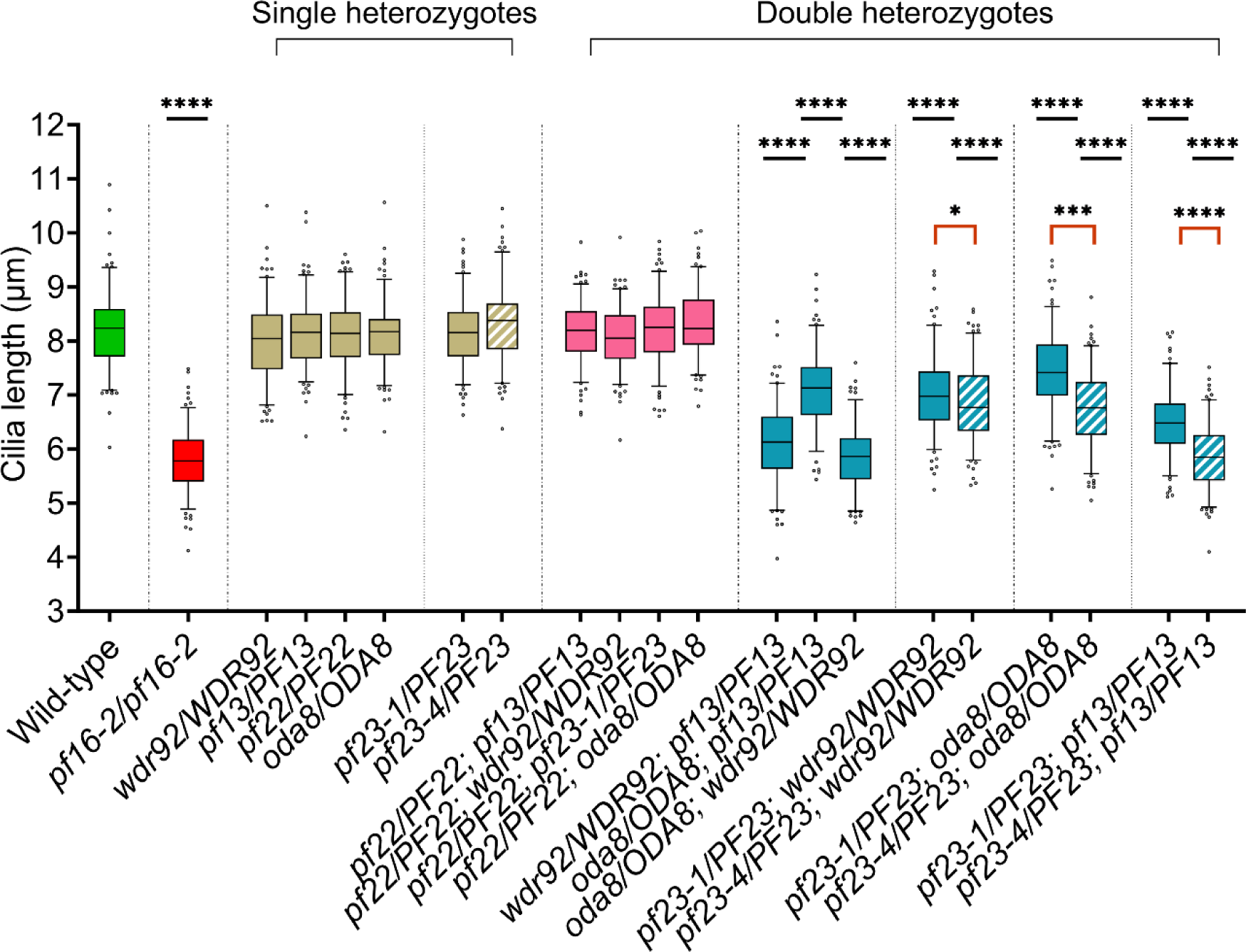
Cilia length of gametic diploids after regeneration for 180 min. Gametic diploids of the indicated genotypes were deciliated and allowed to regenerate for 180 min in 10 μg/ml cycloheximide. Three biological replicates of 50 individual cilia from 50 cells were collected for each strain. The cilia length of each strain is compared to the wild-type diploid. Solid green fill indicates wild-type; single heterozygotes are represented by various solid fill colors. Striped fill indicates double heterozygotes while checkered fill indicates a double heterozygote generated using a null allele of *pf23* (*pf23-4*). Comparison done using a one-way ANOVA with corrections for multiple testing. Red brackets highlight statistical comparison between double heterozygous diploid with either the *pf23-1* allele or *pf23-4* null allele. ‘*’ p < 0.05; ‘**’ p < 0.001; ‘*** p < 0.0001; ‘****’ p < 0.00001; ‘ns’ not significant.

**Table 2:** Mass spectrometry of axonemes from the wild-type strain 4-2P and *pf23-4; cnk11-3*. The relative abundance of peptides from axonemes of *pf23-4*; *cnk11-3* strains normalized to wild-type axonemes is shown. Numbers were derived from conversion of Log2 values. A hyphen (-) or red shading indicates that the peptide was undetectable or extremely low. Proteins with brown shading were decreased to less than 6%, while proteins with tan shading were detected at levels between 20-44%. A hyphen (-) Indicates less than 1/1000. Not detectable. The names in parentheses refer to the associated *Chlamydomonas* mutant or the axonemal complex where it is located. (RSS: radial spoke stand-in, CA: central apparatus; MIP: Microtubule inner protein).

| Protein | Normalized<br>*pf23-4; cnk11-3* | Protein | Normalized<br>*pf23-4; cnk11-3* |
| --- | --- | --- | --- |
| **Outer arms** |  | **ODA docking proteins** |  |
| ODA2 | - | ODA1 | 0.435 |
| ODA4 | - | ODA3 | 0.435 |
| ODA11 | - | ODA14 | 0.233 |
| ODA6 | - |  |  |
| ODA9 | - | **Tubulins** |  |
| | | Tubulin $\beta$ | 1.1 |
| | | Tubulin $\alpha$ | 0.96 |
| **Inner arms** |  | **Dynein arm associated** |  |
| PF9 | - | FAP44 (I1/f tether) | 0.10 |
| DHC10 | - | FAP244 (I1/f tether) | 0.0027 |
| BOP5 | - | FAP43 (I1/f tether) | 1.37 |
| IC97 | - | FAP73 (MIA2) | 0.84 |
| LC7 | - | FAP100 (MIA1) | 0.61 |
| FAP120 | - | FAP57 (BOP2) | 0.87 |
| DHC2 | 0.002 | FAP61 (RS2) | 1.86 |
| DHC3 | - | FAP91 (RS2) | 1.25 |
| DHC4 | - | FAP184 (RSS) | 0.93 |
| DHC5 | 0.0027 | FAP263 (RSS) | 1.09 |
| DHC6 | 0.0027 | DRC4 (PF2) | 0.71 |
| DHC7 | - | DRC1 (PF3) | 0.61 |
| DHC8 | 0.0018 | CCDC146 (MBO2) | 1.13 |
| DHC9 | - |  |  |
| DHC11 | - | **Other structures** |  |
| IDA7 | - | Hydin (CA) | 1.29 |
| IDA4 | - | CPC1 (CA) | 1.5 |
| MOT24 | - | PF16 (CA) | 0.91 |
|  |  | PARCG (IJ) | 0.83 |
|  |  | FAP166 (MIP) | 0.47 |
|  |  | RIB72 (MIP) | 1.14 |
|  |  | RIB43a (MIP) | 0.76 |
| **Light chains** |  |  |  |
| DLC7b | - |  |  |
| TCTEX1 | 0.018 |  |  |
| TCTEX2 | 0.032 |  |  |
| ODA12 | 0.064 |  |  |
| ODA13 | 0.011 |  |  |

Immunoblot analysis of cilia from *pf23-4; cnk11-3* confirm that IC138 and IC2 are missing from *pf23-4*, and the N-DRC component DRC2 and microtubule inner protein RIB43a are unchanged. Immunofluorescence analysis also shows that IC2 fails to localize to the cilia in the *pf23-4* null strain (Fig 8G). These data confirm that the *pf23-4* null mutation shows a more severe assembly defect than *pf23-1*. ODAs and IDAS are affected but other structural proteins are not.

### Preassembly factor PF23 is involved in IC138 modification

Immunoblots of whole cell extracts of *pf23-1* and *pf23-4* haploid strains with an antibody against the IDA I1/f intermediate chain, IC138, shows that two bands are present in whole cell extracts instead of the single band found in wild-type strains. Both bands migrate faster than the band in wild-type (Fig 8C). We transformed *pf23-1* with a wild-type copy of *PF23*. In two independent rescued strains containing a fluorescent tag and FLAG tag, the IC138 protein is restored to the wild-type pattern (Fig 8D, 8E). Since there is no IC138 present in the proteomic analysis of *pf23-4; cnk11-3* cilia, the changes we observe occur exclusively in the cytoplasm of the *pf23-4* mutants (Fig 8F, 8G).

### The severity of the SSNC phenotype in dynein arm preassembly double heterozygotes is gene-specific

Next, we wanted to confirm the SSNC phenotypes observed in the regeneration assay (Fig 7). We repeated the cilia regeneration assay in the presence of cycloheximide for a select group of single and double heterozygous strains and extended the regeneration time to 180 mins (Fig 9). We also wanted to determine whether the phenotype is rescued with increasing time (Fig 9, Supplemental Table 7). As expected, the wild-type diploid strain regrew cilia to approximately 8.19 +/- 0.70 microns (Fig 9, column A), and additional time did not result in longer cilia. The preassembly factor single heterozygous diploids regrew wild-type length cilia. This confirms that these mutations are recessive in this assay (Fig 9, brown shading). We observed that double heterozygous diploids with the *pf22* mutation regenerated cilia with lengths comparable to wild-type (Fig 9, pink shading). Other double heterozygous diploids with combinations that include *oda8, wdr92,* and *pf13*, show different cilia regeneration lengths (Fig 9, aqua shading).

### The *PF23* gene shows allele-specific SSNC

To assess whether the *pf23-1* and *pf23-4* alleles show allele-specific SSNC, we generated double heterozygous diploids using the *pf23-4* allele in combination with the three mutations that produced SSNC in our matrix with *pf23-1* (*wdr92, oda8, and pf13*). We subjected these six double heterozygous diploids to the deciliation and regeneration assay described above. Across all six genotypes, the double heterozygous diploids with the *pf23-4* allele regenerated shorter cilia than those with the *pf23-1* allele (Fig 8D). Since the *pf23-1* allele allows more assembly of ODAs, this may result in a less severe phenotype [73].

### SSNC between *pf23* mutations and other preassembly/maturation mutations occurs through dosage-dependent effects of PF23

To further investigate the mechanism behind the SSNC phenotypes, we utilized PF23 antibody to ask whether PF23 protein abundance is affected by heterozygosity. First, we examined the effect of gene dosage in steady-state gametic cells with the *pf23-4* null mutation in the absence of cycloheximide. Whole cell extracts of wild type diploids (CC-124; CC-5908), double heterozygote *pf23-4/PF23; wdr92/WDR92,* and single heterozygotes *pf23-4/PF23,* and *wdr92/WDR92* were prepared. Immunoblots show that wild-type levels of PF23 protein are produced in the *wdr92/WDR92* single heterozygote and the *oda8/ODA8* single heterozygote but approximately one-half of wild-type levels of PF23 are observed in the *pf23-4* single heterozygote. PF23 abundance is not further reduced in *pf23-4/PF23; wdr92/WDR92* (Fig 10A, 10D) or *pf23-4/PF23; oda8/ODA8* diploid strains (Fig 10B, 10E). PF23 is present at wild-type abundance in the *oda8/ODA8*; *wdr92/WDR92* double heterozygote (Fig 10C, 10F). The presence of the *wdr92* or *oda8* mutations do not affect the level of PF23 protein.

**Fig 10:**
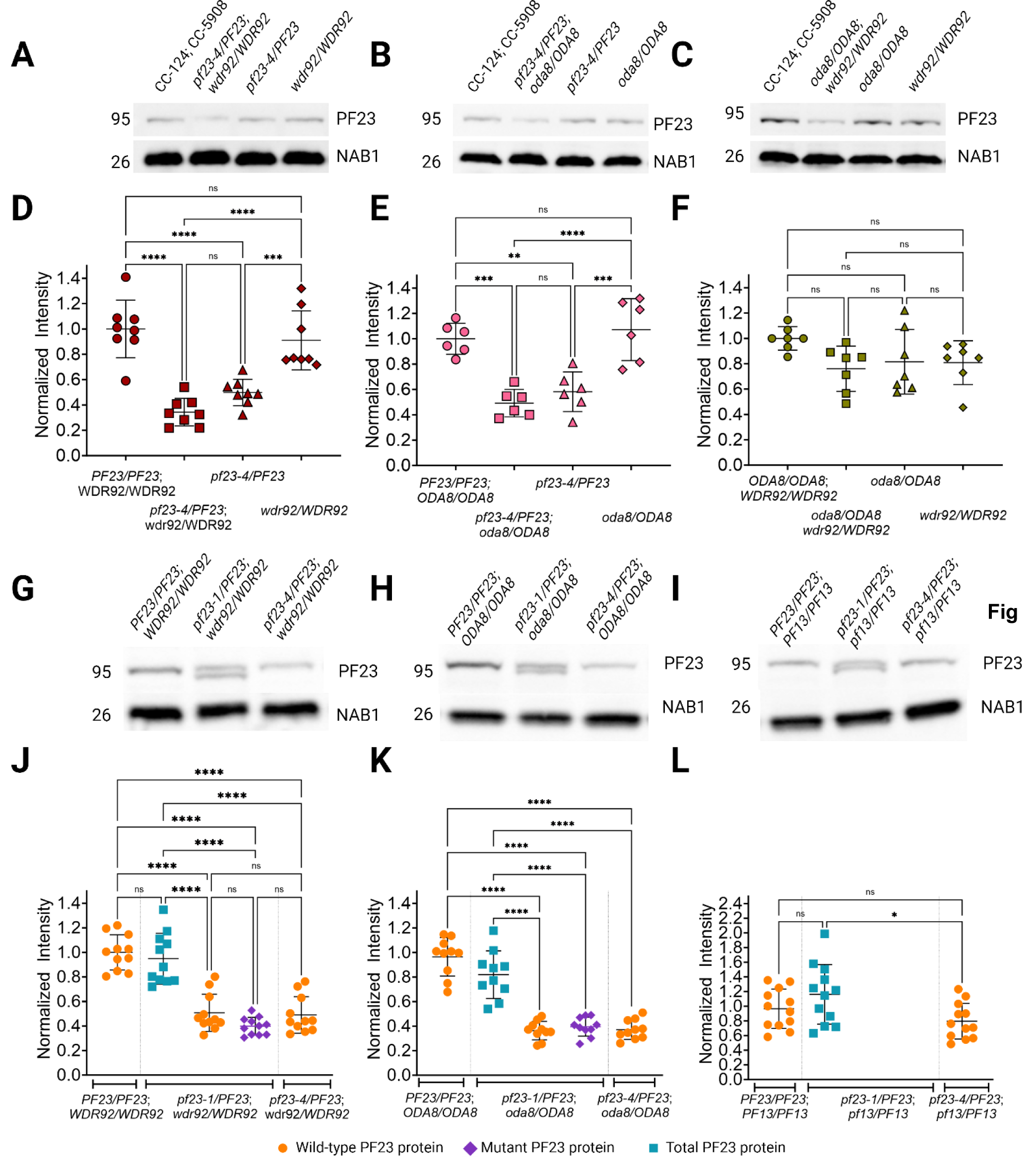
PF23 dynein arm preassembly factor shows a dosage-dependent phenotype in heterozygous strains. (A,B,C) Immunoblots of whole cell extracts of gametic diploid strains of the indicated genotypes. (D,E,F) Quantification of samples in A, B and C. (G, H, I) Immunoblots of cytoplasmic extracts from gametic diploid strains that were deciliated and allowed to regenerate cilia for 60 min in 10 µg/ml cycloheximide. (J, K, L) Quantification of samples in G, H, and I. A total of 50 µg of protein was loaded in each lane. Lane intensities were quantified and normalized to total protein stain. Samples were normalized to the mean of wild type diploids (CC-124; CC-5908). Statistical analysis was performed using a one-way ANOVA. ‘*’ p < 0.05; ‘**’ p < 0.001; ‘*** p < 0.0001; ‘****’ p < 0.00001; ‘ns’ not significant.

Next we assessed the effect of gene dosage and allele specificity of *pf23* in the absence of protein synthesis. In these experiments, we asked if PF23 abundance was regulated by other genes identified in our SSNC screen, as well as the contributions of the *pf23-1* allele. Diploids were deciliated and allowed to regenerate in cycloheximide for 60 mins. These cells were used to prepare whole cell extracts. We compared immunoblots of *pf23-1/PF23*; *wdr92/WDR92* diploid to *pf23-4/PF23; wdr92/WDR92*. As expected, the *pf23-4/PF23; wdr92/WDR92* strain contained roughly half the quantity of PF23 compared to wild-type extracts. In the *pf23-1/PF23*; *wdr92/WDR92* diploid strain, each allele (*pf23-1* and *PF23*) contributes equal amounts of PF23 protein; the total was comparable to that of the wild-type diploid (Fig 10G, 10J). This was also observed when we compared diploids heterozygous for *pf23* and *oda8* (Fig 10H, 10K). When we performed the same assay in diploids heterozygous for *pf23* and *pf13*, PF23 protein abundance was comparable to wild-type in *pf23-4/PF23; pf13/PF13* and the total PF23 protein in *pf23-1/PF23; pf13/PF13* was slightly higher than wild-type, although it did not reach statistical significance (Fig 10I, 10L). We were unable to separately quantify the PF23 protein contributions from the *pf23-1* and *PF23* alleles since the bands did not resolve sufficiently. However, it appears that wild-type PF23 protein is increased in the background of the *pf13* mutation.

## Discussion

### The cytoplasmic pool of axonemal proteins determines cilia regeneration capacity of haploids and diploids

*Chlamydomonas* cells contain a pool of proteins for building cilia that is sufficient to assemble half-length cilia after deciliation in haploid cells if no new protein synthesis occurs [7]. We found that regenerated cilia of wild-type diploids are longer than haploids under these conditions; this is likely to be a direct consequence of the difference of cytoplasmic pool sizes in haploid and diploid cells. It is this limiting pool of proteins in the presence of cycloheximide that provides the sensitized background for our screen. With the limiting pool, we were able to assay the effects of dosage in the double heterozygotes.

Having twice the genetic material of haploid cells is likely to lead to an increase in biosynthetic capacity in diploid cells [101]. In *Saccharomyces cerevisiae*, diploid cells are larger, and have double the number of ribosomes compared to haploid cells [109]. Although this has not been measured in *Chlamydomonas* the same principle may apply. In addition to assembling longer cilia when protein synthesis is blocked, our results show that the rate of cilia regeneration in wild-type diploids is faster than in wild-type haploids. Since both the existing pool and new protein synthesis is required to regrow full-length cilia, we conclude that the faster rate of regeneration observed in these diploids must be a direct consequence of their larger pool size. Interestingly, diploid cells remain able to regulate the total ciliary length even when cilia assembly is faster than in haploids. Cilia length is tightly regulated in *Chlamydomonas* [26]. Changes in cilia length in mutants have not been associated with changes in cell size [110–114]. Diploids supports the idea that cilia length is independent of cell and pool size.

The haploid motility mutants we tested in our screen have a wide range of phenotypes (Fig 2). We anticipated that some double heterozygous diploids would show SSNC in steady-state cells with no stress. However, we observed no obvious motility phenotypes in any of the diploid strains. One explanation is that when the cells are grown under vegetative conditions for two days, there is enough time for them to synthesize the components necessary to build functional cilia. In our matrix, we tested 12 structural ciliary proteins. None of the mutants in these genes show cilia regeneration defects when protein synthesis is inhibited, which suggests that these structural proteins are not the limiting protein(s).

### SSNC among *Chlamydomonas* IFT mutants likely occurs through both poison and dosage interactions

The IFT complexes are composed of 22 proteins [65]. Missense, in-frame deletion, and splice-site mutations that create in-frame exon/intron skipping or inclusion can result in the production of proteins that have lost their normal function but can be poisonous by interfering with assembly or function as occurs between the α- and β- tubulin mutant alleles in *Drosophila* as mentioned previously [59,60].

Five of the *Chlamydomonas* IFT mutants we tested contain missense mutations; they are *IFT144/fla15*, *IFT172/ fla11-; KIF3A/ fla10*-*1, KIF3B/fla8-3,* and *DYHC2/fla24*-*1*. The *IFT81/fla9* allele occurs at a splice-site, while *IFT139/fla17* [66] allele is an in-frame deletion. At the permissive temperature, the haploid strains show mild phenotypes; they assemble cilia and swim, but velocities and numbers of the IFT trains are altered [115].

The strains with null alleles caused by a nonsense mutation (*ift80)* or an insertion (*IFT52*/*bld1-1*) completely lack cilia at all temperatures. All the conditional mutants lose cilia at the restrictive temperature. This phenotype is mirrored in the double heterozygous strains at the restrictive temperature. We hypothesize that the increase in temperature makes the proteins harboring these mutations less stable and they are subsequently degraded. For instance, the *fla24* mutant produces very little FLA24 protein at the permissive temperature; the protein completely disappears at the restrictive temperature, making it null [92]. In this case, even though wild-type IFT complexes can be assembled, the quantity is insufficient to maintain the cilia. It is also possible that incomplete, less stable IFT complexes are formed that lack one or both components missing in the double heterozygote. These incomplete complexes may fall apart at the higher temperature, leaving the cell unable to transport the cargo.

The IFTB mutant *ift72* fails to complement IFTA mutants. IFTB and IFTA function in different phases of ciliary transport. However, IFT172 is unique because it functions in cargo turnover from anterograde to retrograde at the ciliary tip with the microtubule end-binding protein EB1 [87]. In addition, the retrograde defect observed in the *ift72* mutant resembles the IFTA mutants [66,87]. It is possible that the SSNC we observe between IFT172 and the IFTA complex proteins is because IFT172 links retrograde and anterograde transport. A reduction of anterograde IFT together with decreased turnover at the tip facilitated by IFT72 could explain the molecular basis for the SSNC observed in heterozygous diploids with *ift72* and IFTA mutations.

### The *pf23-1* deletion and *pf23-4* null alleles provide new insights into ODA and IDA preassembly in *Chlamydomonas*

The *pf23-1* and *pf23-4* strains show a major difference in assembly of outer dynein arms (Table 2 vs. Yamamoto *et.al.* [73]). We propose that the DYX domain, which is deleted in *pf23-1*, is responsible for IDA assembly. Since ODAs are reduced around 50% in the *pf23-1* strain, it is likely that the DYX domain and the remaining protein works cooperatively to properly assemble ODAs (Table 2). Studies of DYX1C1/PF23 mutations in mice, zebrafish and humans that are presumed to be null show loss of both ODAs and IDAs from the cilia [41,44]. *Chlamydomonas* PF23 was suggested to only have a role in assembly of IDAs based on studies of the *pf23-1* allele. The isolation and characterization of a null *pf23* allele in *Chlamydomonas* shows that PF23 is responsible for assembly of both ODAs and IDAs, as in other organisms. Equal amounts of wild-type and mutant protein are found in the *pf23-1/PF23* heterozygote.

This indicates that while the DYX domain is important for PF23 function, it is not necessary for PF23 stability. Previous studies show that phosphorylation is important for IC138 function in cilia [22,113], and requires casein kinase 1 [116]. We observed altered mobility of IC138 in whole cell extracts of *pf23* mutants. Since IC138 is completely missing from isolated cilia based on our mass spectrometry and immunoblot data (Fig 7), the difference in size must arise from modifications that occur in the cytoplasm. We suggest that IC138 undergoes two independent modification events; one in the cytoplasm that is dependent on PF23, and another in the axoneme that requires casein kinase 1. We hypothesize that the cytoplasmic modification of IC138 is necessary for its assembly into the I1/f IDA dynein, or to facilitate its transport into the axoneme by IFT through interactions with its adapter IDA3 [52]. Further experiments are needed to determine the nature of the modifications. It is not known if other assembly factor mutants cause similar changes to IC138. Previous studies showed that ODAs require factors (ODA5, 8, and 10) that help them assemble efficiently and make them competent for axonemal transport and/or binding. PF23 and other assembly factors may serve a similar role for IDAs.

### The abundance and stability of cytoplasmic ODAs and IDAs proteins differ among the DNAAF mutants

In the *pf22* mutant cytoplasm, the ODA heavy chains are present at wild-type levels, but they fail to assemble. This suggests that while PF22 is important for ODA preassembly, it is not involved in ODA heavy chain stability [69] and PF22 is not known to interact with chaperone proteins as observed for other DNAAFs. In contrast, all three ODA heavy chains (α,β, and γ) and one IDA subunit are depleted in *pf13* cytoplasm, and the remainder of ODA heavy chains fail to assemble with the intermediate chains. Therefore, PF13 is necessary for both stability and assembly of ODAs [71]. ODA and IDA subunits are also reduced or absent in the cytoplasm of *pf23* and *wdr92* mutants, highlighting a role in dynein stability [41,42,57,72,73].

All three DNAAFs identified in our screen interact with other DNAAFS and chaperone proteins. The canonical R2TP chaperone complex is comprised of preassembly factors RUVBL1 (Pontin), RUVBL2 (Reptin), RPAP3 (RNA polymerase III associated protein), and PIH1D1/MOT48/IDA10 with a PIH-domain (Protein Interacting with HSP90) [43]. The R2TP complex is widely involved in the binding and folding of nascent proteins [43,46,117] In studies of human airway epithelial cells, PF13 was shown to form an early preassembly complex with DNAAFs HEATR2, a heat-repeat protein, and SPAG1, a protein with an RNA-polymerase II-associated domain [118]. Both PF13 and SPAG1 form a non-canonical R2TP complex with RUVBL1 and RUVBL2 [45,46]. Preassembly factor PF23 interacts with multiple DNAAFs and chaperone proteins that include PF13, MOT48/PIH1D1, PIH1D3/TWISTER/TWI1, HSP90 and the TriC tubulin chaperone complex [44,119]. WDR92 is associated with RPAP3, SPAG1, and the prefoldin complex [42,46,120]. Prefoldin functions as a monitor for misfolded proteins [121–123].

Unlike most DNAAFs, only ODAs are affected in the *oda8* mutant. Wild-type levels of ODA heavy chains α,β, γ and IC2 are found in the *oda8* cytoplasm. The *oda8* mutant, like *pf13,* fails to assemble ODA intermediate chains with ODA heavy chains but this interaction is not important for ODA subunit stability [124]. ODA8 has an additional role in preparing ODAs for transport into the cilia, since transport of IC2 into the cilia is very infrequent in the mutant [49].

In double heterozygous diploids *pf23-4/PF23-4; wdr92/WDR92* and *pf23-4/PF23- 4; oda8/*ODA8, one-half of the wild-type levels of PF23 are present. This suggests that PF23 transcription or translation is not increased in these strains to counteract the deficit from the *pf23* null allele, and PF23 stability is not dependent on WDR92 or ODA8. In the diploids heterozygous for *pf13* and either *pf23-1* or *pf23-4* mutations, PF23 protein abundance is closer to wild-type levels in the presence of cycloheximide. This observation suggests that a change in dosage of PF13 regulates the amount of PF23 in the cytoplasm. How this regulation occurs is unknown.

### Dynein arm preassembly factors show dosage effects through reduction of multiple pathway components

In our screen, cells must regenerate cilia using the existing cytoplasmic pool. In a wild-type diploid, both alleles of a *DNAAF* gene contribute to the protein pool, and all the dynein components are stable and available for assembly. We did not detect SSNC in any diploids heterozygous for *pf22*. One explanation for this is that PF22 has a different role in dynein preassembly than other DNAAFs. In the diploids that show SSNC, most have a combination of two DNAAFs (PF13, PF23, AND WDR92) that affect the stability of the dynein motors. The SSNC we observe with in ODA8 could be due to the combinatorial effects of inefficient dynein preassembly and reduced production of mature dyneins that can be transported into and bind the axoneme.

The strains *pf13-1/PF13; oda6/ODA6,* and *pf13-1/PF13; oda9/ODA9* are heterozygous for mutations in one dynein preassembly factor and one structural protein. The ODA6 and ODA9 proteins (IC2 and IC1) stabilize each other [79]. IC2 is completely missing in the *oda6* mutant, and severely reduced in the *oda9* strain [80]. It is possible that the SSNC we observe in these two strains is due to the reduction of three proteins. First, PF13 protein is reduced by half because it is heterozygous; there is less available to participate in dynein preassembly and the total pool of ODA heavy chains could be reduced since PF13 stabilizes them. Second, ODA6 or ODA9 are reduced by half; the pool of intermediate chains may be limited. Together, the reduction of PF13 available for preassembly and a decrease in dynein arm precursors could explain the impaired ability of these two heterozygotes to regenerate their cilia.

In the *pf23-4/PF23*; *wdr92/WDR92* double heterozygote, PF23 is reduced by half based on our immunoblot data. This is true whether or not protein synthesis is inhibited. We hypothesize that WDR92 is also reduced by half, although we were unable to test this assumption. If we assume that this is true, in the *pf23-4/PF23*; *wdr92/*WDR92 strain ODA and IDA protein stability is compromised and there are fewer subunits available for regeneration.

In *Xenopus*, live imaging of the cytoplasm of multiciliated cells (MCCs) shows that dynein preassembly factors and dynein subunits colocalize in foci called dynein assembly particles (DyNAPs) [125]. These foci are dynamic; FRAP (fluorescence recovery after photobleaching) imaging shows that the dynein subunits exist stably within the particles, while about 50-80% of the DNAAFs move in and out of DyNAPs [125]. Colocalization imaging in the *Xenopus* MCC cytoplasm shows that PF13/Ktu colocalizes with 5 different preassembly factors (ZMYND10, PIH1D3, SPAG1, LRRC6 and PF23/DYX1C1) and three chaperones (RUVBL1, RUVBL2, and HSP90) in DyNAPs [125]. In another study, co-immunoprecipitation and yeast two-hybrid assays identified that FBB18 interacts with these same three chaperones, as well as PF22, WDR92, ODA7 and PF23/DYX1C1, but not SPAG1, MOT48, RPAP3, or ZMYND10 [42,126]. PF13 and PF23 interact by immunoprecipitation [44]. This suggests that there are DyNAPs with different compositions. Knockdown of HEATR2 in *Xenopus* resulted in fewer DyNAPs; and the movement of PF13/Ktu in DyNAPs decreased substantially [127]. It is possible that loss of other DNAAFs will decrease the number of DyNAPs, but this has never been examined. Another observation is that ODA and IDA subunits occupy different compartments within the same DyNAP [127]; it is likely that different combinations of DNAAFs could be recruited to these sub-compartments to assemble ODAs and IDAs, but this was not tested. From these data, we surmise that multiple configurations of DyNAPs can be formed that contain both shared and unique combinations of DNAAFs that participate in different stages of dynein preassembly. Previous data from *wdr92* mutant studies show that a high molecular weight complex with PF23 disassembles in the *wdr92* mutant. This supports the idea that PF23 and WDR92 function together in a hub for preassembly [42,57], and that WDR92 may affect DyNAP formation in a similar way to HEATR2. Because PF23 interacts with multiple DNAAFS, we hypothesize that the pool of PF23 proteins is shared between different preassembly complexes (DyNAPs) with different roles in dynein assembly. In strains heterozygous for PF23, it is likely that there are fewer preassembly complexes (DyNAPs) with PF23 and WDR92. We predict that other PF23-containing DyNAPs will also be reduced and disrupt different aspects of dynein preassembly.

Having half the quantity of PF23 is enough for normal cilia regeneration since we do not observe phenotypes in *pf23/PF23* single heterozygotes. However, when this PF23 deficit is combined with a reduction in another preassembly factor, the collective effect of reduced preassembly machinery and disruption of other DyNAPs, together with a deficit in dynein heavy chains compromises the cell’s ability to produce enough dyneins to rebuild the cilia. We were limited to testing SSNC among pre-existing mutant strains available through the *Chlamydomonas* Resource Center. Although there are currently 17 known DNAAFs, we were only able to obtain mutants in 5 of them; and except for *pf23-1*, they are all null alleles (Table 1). The tools for generating new alleles and tagged strains have improved since we initiated this project. In the future, it will be interesting to test whether combinations of DNAAFs and chaperone proteins show SSNC.

### Can double heterozygosity in motile cilia genes play a role in human disease?

In humans, pathogenic variants in 58 genes [128] cause a rare disease called primary ciliary dyskinesia (PCD), characterized by recurrent lung infections, neonatal respiratory distress, bronchiectasis, otitis media, situs inversus/ambiguus, and male infertility [2,129,130]. Reports of SSNC or digenic inheritance (DI) with motile cilia genes are rare. One study identified patients with heterozygous variants in *DNAH6*, an IDA heavy chain gene, and another variant in *DNAI1*, an ODA intermediate chain gene. These individuals had heterotaxy, a PCD-associated phenotype [124]. Knockdown of *DNAH6* in mice heterozygous for *dnaI1* resulted in motile cilia defects, which suggests that heterozygous *dnah6* and *dnai1* mutations are responsible for the patient phenotypes [131]. Roughly 30% of patients with a clinical PCD diagnosis have unknown genetic etiology despite genetic testing and/or exon sequencing. A small subset of patients with a PCD diagnosis are heterozygous for variants in one or more PCD genes [132–135]. This implies that either the correct causative gene was missed due to a complex variant, a variant in a PCD gene not included in the test set, a variant is in a non-coding region, or the second hit is in a different gene than the one already identified [135–139]. Our results reinforce the findings that PCD mutants show a recessive inheritance pattern. Although our steady-state double heterozygous diploids do not show obvious motility or cilia defects, the possibility of SSNC should be considered when looking at human PCD patients. Individuals with milder PCD phenotypes might be candidates for SSNC. Double heterozygosity phenotypes may be exacerbated under stress conditions that include viral infections and other environmental exposures and produce PCD-like phenotypes. The implications of this work are that individuals who are double heterozygotes for two motile cilia genes may encounter difficulties with recovering from assaults that damage their cilia.

## Materials and Methods

### Strains and Diploid Generation

The strains used in this study are listed in Table 1. To generate diploid strains, single mutant haploid strains wild-type for *AC17* were crossed with strains CC-5908 or CC- 5909 with the genotype *ac17; TG-aphviii* (Supplemental Table 1) that are mating-type plus or mating-type minus, respectively. Mutant strains with reciprocal selectable genotypes were mated and then plated onto acetate-free plates supplemented with 10 µg/ml paromomycin. Plates were incubated in constant light at 25°C for 3-5 days until colonies appeared. Each diploid colony was screened by PCR for heterozygosity at the mating-type locus before further analysis. Available PCR markers were used to genotype strains when appropriate. Primers are listed in Supplemental Table S1. Strain CC-5485 [103] with *aphviii* insertions was used to obtain strains CC-5908 and CC-5909. The *aphviii* insertion in CC-5485 does not cause a motility phenotype [103]. The *oda11* strain CC-2672 from the *Chlamydomonas* Resource Center has a second hit that results in short cilia. This strain was backcrossed to wild-type to remove it.

### Deciliation, regeneration, and imaging

For deciliation and regeneration assays, cells were plated onto R (rich) medium plates for two days at 25^°^C [140,141]. They were transferred into 1 ml of liquid R medium in a 1.5 ml Eppendorf tube and rocked under light for 3 hrs to allow cilia to assemble. The cells were placed on ice, spun at 10,000 x g for 30 sec and resuspended in 1 ml cold deciliation solution (10 mM Tris pH 7.5, 150 mM D-mannitol and 1mM CaCl_2_). pH shock was performed with 30 µl 0.5N acetic acid followed by 30 µl 0.5M sodium hydroxide [52] and then checked for deciliation by phase microscopy with a 40x objective. Cells were spun and resuspended in 500 µl R medium with or without 10 µg/ml cycloheximide (Sigma Aldrich) and allowed to regenerate cilia for the times specified. A 40 min timepoint was selected for most experiments because it allowed screening to be completed in a reasonable amount of time. Cells were fixed with a 1:10 dilution of 2% glutaraldehyde (Sigma Aldrich) in 0.2 M phosphate buffer pH 7.5 (0.31 g NaH_2_PO4 and 1.09 g Na_2_HPO4 in 50 ml water). Cells were then allowed to adhere to glass slides coated with poly-L-Lysine solution (Sigma Aldrich). Excess cell volume was aspirated, and the slides were air-dried before imaging. For initial diploid screening of cycloheximide treated cells, fixed samples were imaged using phase contrast microscopy on a Zeiss AxioPhot with a 20x Neofluar objective lens and a NA of 2.0 (Carl Zeiss AG, Oberkochen, Germany). Images were captured using a Phantom Miro eX2 camera and Phantom Camera Control Application 2.6 (Vision Research, Wayne, NJ) using 640 x 480 resolution. Cilia length was quantified using ImageJ. For immunofluorescence staining, protocol was described previously [107]. Primary antibody was anti-acetylated α-tubulin mouse monoclonal clone 6-11B-1 (1:500 dilution, Sigma, T7451-200, lot #077M4751V). Secondary antibody was Alexa Fluor 488 donkey anti-mouse IgG (1:1,000 dilution, Invitrogen, A21202, lot #2266877). Images were false color stained using ImageJ.

### Genome editing

CRISPR genome editing protocol and associated reagents are described [107].

### Axoneme isolation and mass spectrometry

Axonemes of wild-type strain WT 4-2P (generated from a cross between CC-124 and CC-125), strains *pf23-4; cnk11-3* (2-3, 3-1 and 5-1), and *cnk11-3* (1-7, 1-4, and 5-1) [110] were prepared as sets of three biological replicates using the dibucaine method [142]. 100 μg of protein from each sample were submitted for tandem mass spectrometry analysis to the Danforth Plant Science Center St. Louis, MO [143].

### Immunoblotting

Immunoblotting was performed using gametic whole cell extracts. Protein from the cytoplasm and the cilia are included in the extract. Each strain was grown on one TAP plate for 2 days at 25°C, then placed at room temperature for 3 days to allow cells to become gametic. Each plate was resuspended in 10 ml of M-N/5 and rocked in light at room temperature for 3 hr. Cells were spun at 2,000 x g for 5 min, resuspended in 10 ml of autolysin [144] for 1 hr. Next cells were spun at 2,000 x g for 5 min and resuspended in 600-750 μl of cell lysis buffer (20 mM Tris–HCl pH 7.5, 5 mM MgCl_2_, 300 mM NaCl, 5 mM Dithiothreitol, 0.1% Triton-X100 [93], with 20 μM phenylmethylsulfonyl fluoride (Sigma, Lot # 050M1688) and the Protease Inhibitor Cocktail for plants (Sigma, p9599, lot #086K4075). The cell suspension was passed 10 times through a 22 ½ gauge needle and syringe, or frozen overnight at -80°C to generate cell lysate. The lysate was spun at 21,000 x g for 30 min to remove non-lysed cell bodies, organelles and debris. The supernatant was transferred to a fresh 1.5 ml Eppendorf tube and protein concentration was measured using Bradford assay (Bio-Rad Protein assay Dye reagent Concentrate (#5000006, lot 64227541). Samples were frozen at -80°C. For protein electrophoresis, 5x sample buffer (3.78 g Tris, 5g SDS, 25 g sucrose, in 90 ml, pH to 6.8, 0.04 g bromophenol blue in a final volume of 100 ml) with freshly added β-mercaptoethanol to a final concentration of 5% was added to the sample at 1X concentration and heated in a dry bath for 10 min at 70°C. 50 μg of each sample was loaded onto a 4-20% Novex Tris-Glycine gel (XP04205BOX) and run at 225 volts for 45 min. Following transfer onto Immobilon-P 0.45μm PVDF membranes (Millipore Sigma), each membrane was incubated with Invitrogen™ No-Stain™ Protein Labeling Reagent (A44717) according to manufacturer’s instructions. Subsequent blotting steps followed standard procedures [68]. For antibody probing, the membranes were cut horizontally at the appropriate size marker for the protein to be examined. Each section of membrane was incubated with its respective primary and secondary antibody sequentially. Primary antibodies used were rabbit anti-PF23 CT299 [57] (1:1000, a gift from Dr. Stephen King), rabbit anti-IC138 (1:2000, a gift from Drs. Winfield Sale and Lea Alford), rabbit anti-RIB 43a (1:2000, a gift from Dr. Mary Porter), rabbit anti-NAB1 (1:10,000, Agrisera, catalog #AS08 333, lot #1311), and monoclonal mouse anti-IC2 clone 1869A (1:10,000, Sigma, D-6168, lot #92H4862). Secondary antibodies used were goat anti-rabbit HRP (Sigma-Aldrich, A6154-1ML, SLCL2476) at 1:5000, 1:5000, 1:10,000, and 1:10,000 dilutions respectively. Goat anti-mouse HRP (catalog 82-8520, lot #WD321502) was used at 1:10,000 dilution. Membranes were visualized using the Invitrogen iBright 1500 Imager with the universal channel to detect total protein stain and chemiluminescence.

### Recombineering and rescue of *pf13-1* and *pf23-1*

To rescue the *pf13-1* strain, we utilized the recombineering approach detailed in Mackinder *et.al.* [93]. The BAC strain 23P6 containing the *PF13* gene (Cre09.g411450, Phytozome v5.6) was obtained from the *Chlamydomonas* BAC library (http://chlamycollection.org) [145]. Recombineering construct PLM099 (Supplemental Fig 3A) that contains the Venus-YFP-FLAG tag and *aphviii* paromomycin resistance cassette was chosen for recombineering with 23P6. Following recombineering, two colonies were selected for further analysis after running isolated plasmid DNA on a 0.8% agarose gel and checking for the expected size of 10 kb (Supplemental Fig 2B). The two colonies were then digested with restriction enzymes *Nde*I and I-*Sce*I to confirm the presence of the PLM099 backbone and the *PF13* gene (Supplemental Fig 2C). DNA from *PF13::*YFP-FLAG #2 was isolated from a 4 ml overnight culture was used to transform *pf13-1* as described [107]. PCR to detect the presence of the transgene was performed on a subset of colonies (Supplemental Fig 2D) that showed a rescued phenotype of swimming in liquid 96-well plates under a dissecting microscope (*pf13-1* is non-motile. To identify the transgene, PCR primers were used to amplify the junction between the 3’ end of *PF13* and the 5’ end of the YFP-FLAG tag (Supplemental Fig 2E, Supplemental Table 1). The presence of the tagged protein was identified by immunoblotting using anti-FLAG antibody (F1804 monoclonal antibody, clone M2 in mouse, SLCK5688, Sigma Aldrich, Supplemental Fig 2F). Rescue was confirmed by measuring cilia length, swimming velocity and beat frequency (Supplemental Fig 3G, 3H, 3I). Similar steps were followed for rescue of *pf23-1*.

### RNA and cDNA isolation and analysis

Cells for RNA isolation were collected from R plates grown in bright light for two days at 25°C. RNA was prepared using Trisol Reagent (Ambion, catalog 15596018, lot#265712) and quality was assessed using a 0.8% agarose gel. cDNA was generated using SuperScript IV VILO MasterMix (ThermoFisher) [143]. Primers in Supplemental Table 1 were used to detect cDNA.

### Statistical Analysis

Statistical analyses were performed using GraphPad Prism 9 software. Comparisons were performed using a One-Way ANOVA with corrections for multiple testing unless otherwise indicated.

### Whole genome resequencing and analysis

Whole genome sequencing was performed as described previously [68]. In instances where SnpEff [146,147] was unable to identify the causative mutation, the whole genome resequencing data was manually analyzed using IGV and alignment to reference sequences CC-124 and CC-125.

## Acknowledgements

We thank Drs. Steve Brody, Tim Schedl, Moe Mahjoub, and Zachary Payne for useful comments. We thank Dr. Steve King for the antibody to PF23.

## Supporting Information Table Captions

Table S1: PCR primers generated to genotype strains used in this work. Where indicated, the PCR product is digested with a restriction enzyme to distinguish between wild-type and mutant genotypes. ‘No band in mutant’ indicates the presence of an insertion that is too large to amplify during the extension time needed to amplify the PCR products with the same primers in wild-type. *Designed using the method for allele-specific primers (ASP)[148]. ** The mating-type primers can be run simultaneously to generate two bands that distinguish between both mating-types.

Table S2: Raw Log2 values from TMT Mass Spectrometry

Table S3: SNPEff variant output from whole genome sequencing analysis of haploid mutants

Table S4: Cilia length, swimming velocity, and beat frequency of *Chlamydomonas* haploid mutants

Table S5: Comparison of cilia length, swimming velocity, and beat frequency of *Chlamydomonas* haploids and diploids

Table S6: Cilia length, swimming velocity, and beat frequency of *Chlamydomonas* single and double heterozygous diploids

Table S7: Cilia length of regenerating haploids and diploids +/- cycloheximide

Table S8: Cilia length of regenerating single and double heterozygous diploids 40 mins post-deciliation in cycloheximide

Table S9: Cilia lengths of *pf23* mutants and *pf23* rescue strains

Table S10: Cilia lengths of single and double heterozygsous diploids 180 mins post-deciliation in cycloheximide

Table S11: Quantification of immunoblots for Fig 10 and Fig S6

## Supporting Information Figure Captions

Fig S1: **Whole genome resequencing analysis of the *pf13-1* and *oda4-1* strains.** (A) Reads spanning the *PF13* locus (Cre09.g411450) from the *pf13-1* mutant show a 44.7 kb deletion in the mutant strain compared to wild-type CC-124 and CC-125 strains. The IGV track at the bottom in blue shows the genes in that region (B) The genes and corresponding protein products deleted in *pf13-1* strain as annotated in Phytozome 13 [149]. Numbers indicate the order in which the genes are arranged, starting with *PF13*. (C) PCR confirmation of the regions deleted in *pf13-1* using primers listed in Supplemental Table S1. The green check marks show that the sequence is amplified in both wild-type (CC-124) and *pf13-1* and indicate the regions just outside the deletion breakpoints on either side. A red ‘X’ indicates that the sequence between the primer pair was amplified in wild-type but not in *pf13-1*. (D) IGV snapshot of reads spanning the region of the *oda4-1* (Cre09.g403800) showing a 5 bp deletion in exon 22 out of 30 exons. Regions containing the deletions are indicated with a red arrow and bracket.

Fig S2: The ciliary assembly defect of *pf13-1* is rescued by transformation with a *PF13:YFP-FLAG-TG* construct generated using recombineering cloning. (A) Amplification of the PLM099 recombineering plasmid that contains the YFP-FLAG tag and paromomycin selectable marker with PF13-specific 50 bp homology primers (Supplemental Table S1). (B) DNA isolated from two independent bacterial colonies with PLM099 after recombineering with BAC 23P6 (from *Chlamydomonas* BAC library) containing *PF13*. The top band (labelled with an asterisk) likely represents BAC DNA that did not undergo recombineering. (C) Restriction digest of DNA from the two colonies isolated in (B). I-*Sce*I is used for linearization of the plasmid, while *Nde*I + *Xba*I digest indicates the presence of the PLM099 vector backbone along with the *PF13* gene. (D) PCR was performed on a subset of *Chlamydomonas pf13-1* strains transformed with *PF13:YFP-FLAG* #2 to verify the presence of YFP-FLAG tag using primers within the *Venus-YFP* sequence. A strain with *CAH6:YFP-TG* [150] was used as a positive control. (E) PCR confirmation of fusion of the *YFP-FLAG* tag with the 3’ end of the *PF13* gene in two of the strains (A2 & B8 renamed *pf13-1; PF13*:*YFP-FLAG- TG* #1 and #2) tested in (D). (F) Immunoblot of transformed strains in (E) using an anti-FLAG immunoblot. The expected size of PF13 is 75 kDa. The observed bands are larger due to the YFP and FLAG epitope tags present at the C-terminus of PF13. (G,H,I) Cilia length, swimming velocity and beat frequency are rescued in the strains *pf13-1; PF13*:*YFP-FLAG-TG* #1 and #2. CC-124 and CC-125 are wild-type controls. *pf13 is* non-motile and indicated by an ‘X’.

Fig S3: **Genotyping of diploid strains using PCR markers against selectable markers for *ac17* and *aphviii.*** Haploid strains CC-5908 and CC-5909 carry an insertion in the *ATG17* gene, and a mutation in *ac17*, but are wild-type swimmers. CC-124 and CC-125 are wild-type at both loci. The three diploid strains are heterozygous for the *atg11* and *ac17* mutations. (A) PCR primers for the *ATG17* gene[103] were used to detect the wild-type copy of the gene in CC-124, CC-125 and three diploid wild-type strains (lanes 3A-7A). No amplification is observed in CC-5908 and CC-5909 (lanes 1 & 2), since the *aphviii* paromomycin resistance gene is inserted into the *atg11* gene and is too large to amplify with the condition used. (B) PCR primers were designed to detect the junction between the *ATG11* gene and the *aphviii* insertion in the *ATG11* gene. This junction is present in the CC5908 and CC-5909 strains that carry the insertion (lanes 1B & 2B), as well as the heterozygous diploids (lanes 5B-7B). (C) PCR primers were designed to amplify across the region containing the mutation in the *AC17* gene [151]. Digestion with *Hae*III generates different sized products in the *ac17* containing strains (lanes 1C & 2C) compared to CC-124 and CC-125 wild-types (lanes 3C & 4C). The heterozygous diploid strains (5C-7C) generate both sets of digest products. (D) A map of the *AC17* PCR product digested with *Hae*III.

Fig S4: **Isolation and analysis of wild-type strains transformed with *aphvii* targeted to exon 1 of the *PF23* gene.** (A) PCR screening of 7 transformants from *PF23* CRISPR insertional mutagenesis. One colony, B10 fails to amplify the exon 1 region targeted by the primers and amplifies the *aphvii-PF23* junction in exon 1. (B) Strain H9 was retrieved by screening an additional 11 strains from the same transformation. The insertion is oriented in the forward direction. (C) A second transformation produced strains 2-3 (*pf23-2)* and 4-3 (*pf23-3*), the insertion in both strains is oriented in the reverse direction. (D) cDNA analysis of *PF23* CRISPR transformants to detect whether mRNA is disrupted. A band was amplified in strain B10, and it was excluded from further analysis. Strains 2-3, 4-3 and H9 (*pf23-2*, *pf23-3,* and *pf23-4* respectively) failed to generate a cDNA amplicon for exons 1 and 2.

Fig S5: **Genotyping of *pf23-1; PF23* transgenic rescue strains.** Strain *pf23-1* contains an in-frame deletion in exon 5 and was rescued with wild-type *PF23* constructs tagged with either m-scarlet or neon green. These are listed at the top of the Fig. The right labels indicate the primers used (See Supplemental Table S1). (A) The CC-124 and *pf23-4* strains both amplify wild-type PCR products at exon 5. The *pf23-1* amplicon is smaller due to the in-frame deletion. Both rescue strains carry the smaller *pf23-1* amplicon and a wild-type exon 5 amplicon. (B) Amplicon of the junction between the 3’ end of *PF23* and the 5’ end of m-scarlet in the rescue strain. Asterisks indicate non-specific bands. (C) Amplicon of the junction between the 3’ end of *PF23* and the 5’ end of neon green. (D) Amplicon of the junction between the 3’ end of *PF23* and *aphvii* in the *pf23-4* null strain with an insertion in exon 1. (E) *pf23-4* fails to amplify a band in exon 1 due to the *aphvii* insertion.

Fig S6: **Quantification of PF23 abundance in diploid and haploid strains with different *WDR92* genotypes.** (A) Representative blot with 50 µg protein of the indicated strains. Each strain within the brackets indicates a biological replicate. Haploid strains with genotype *WDR92* included CC-5908 and CC-124. IC2, NAB1 are used as loading controls. Total protein stain was used for quantification. (B) Quantification of samples (biological and technical replicates). All samples were normalized to the mean of the wild-type diploids. A one-way ANOVA was used to assess statistical significance.

## Notes

### Competing Interest Statement

The authors have declared no competing interest.

